# The Autonomic Nervous System Differentiates Between Levels of Motor Intent and Hand Dominance

**DOI:** 10.1101/2020.06.01.128140

**Authors:** Jihye Ryu, Elizabeth Torres

**Affiliations:** Psychology Department, Rutgers University Center for Cognitive Science, Rutgers University, Piscataway, NJ 08854, USA; Psychology Department, Rutgers University Center for Cognitive Science, Computational Biomedicine Imaging and Modeling Center at Computer Science Department, Rutgers University, Piscataway, NJ 08854

**Keywords:** embodied cognition, agency, action ownership, network analysis, motor variability, motor control, voluntary motion, precision medicine

## Abstract

While attempting to bridge motor control and cognitive science, the nascent field of embodied cognition has primarily addressed intended, goal-oriented actions. Less explored however, have been unintended motions. Such movements tend to occur largely beneath awareness, while contributing to the spontaneous control of redundant degrees of freedom across the body in motion. We posit that the consequences of such unintended actions implicitly contribute to our autonomous sense of action ownership and agency. We question whether biorhythmic activities from these motions are separable from those which intentionally occur. Here we find that fluctuations in the biorhythmic activities of the nervous systems can unambiguously differentiate across levels of intent. More important yet, this differentiation is remarkable when we examine the fluctuations in biorhythmic activity from the autonomic nervous systems. We find that when the action is intended, the heart signal leads the body kinematics signals; but when the action segment spontaneously occurs without instructions, the heart signal lags the bodily kinematics signals. We posit that such differentiation within the nervous system, may be necessary to acquire the sense of action ownership, which in turn, contributes to the sense of agency. We discuss our results while considering their potential translational value.

## 1. Introduction

The field of embodied cognition (EC) has provided a powerful theoretical framework amenable to bridge the gap between research probing our mental states and research investigating our physical actions [1–3]. Indeed, within the framework of EC, the construct of agency conceived as a cognitive movement phenomenon [4–6], may provide a way to finally connect the disparate fields of cognitive science and motor control. An important component of agency is action ownership [5, 7, 8], *i.e*. the sense that sensory consequences of the actor’s action are intrinsically part of the actor’s inner sensations. When the actor owns the action, s/he has full control over those sensations that are internally self-generated and self-monitored by the actor’s brain, and yet extrinsically modulated by external sensory goals. A critical aspect of this internal-external loop is the identification of the level of actor’s intent, and its differential contribution to the action’s intended and unintended sensory consequences.

In recent years, a body of knowledge has increased our understanding on the sensory consequences derived from intentional actions, as such action components deliver an overall sense of agency [9, 10]. Less explored however, have been parts of the action that are unintended, or that transpire spontaneously and largely beneath awareness. Such actions’ components exist at the involuntary and at the autonomic levels of neuromotor control (Figure 1). They do not require explicit instructions or precisely defined external goals, yet they too contribute to the differentiation of levels of intent in our actions [11, 12]. More importantly, at the cognitive level of decision making, these unintended movements contribute to the acquisition of decision accuracy, within the context of motor learning induced by different cognitive loads [13, 14].

**Figure 1.**
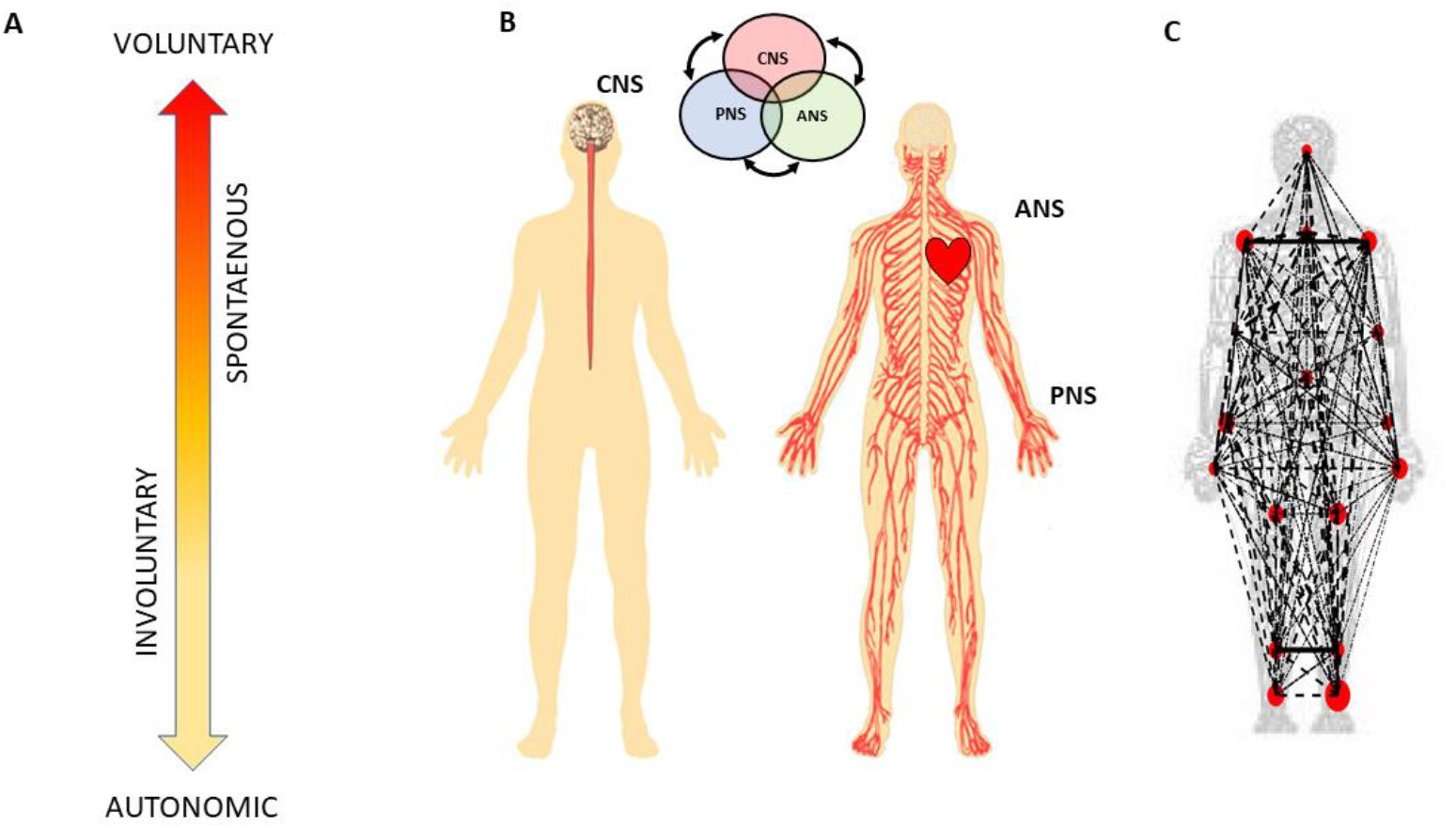
Defining quantitative aspects of agency for the study of embodied cognition. **(A)** Phylogenetically orderly taxonomy of nervous system functions involving different levels of voluntary control (intent) ranging from deliberate to spontaneous movement segments, to involuntary motions and autonomic control. Multi-layered signals contributing from each of these layers are proposed to differentially contribute to the sense of action ownership and to the overall sense of agency via sensory consequences preceded by different levels of intent. **(B)** Contributions of the central and peripheral nervous systems, including the autonomic nervous system (ANS), can be tracked in a closed loop that helps the autonomous realization of intended thoughts into physical actions under volitional control. **(C)** Network connectivity analyses of kinematics and heart biorhythmic signals encompassing these levels of control enable the study of agency through objective quantitative methods.

At the motor control level, autonomous and spontaneous movements are important to develop a sense of action ownership in the face of motor redundancy [15]. They require the coordination of many degrees of freedom (DoF) across the body. Thus, as we produce fluid and timely goal-oriented actions, kinematic synergies self-emerge and dynamically recruit and release the bodily DoFs, according to task demands [16–18]. Conscious decisions generating movements that attain external goals take place as the brain interweaves deliberate and spontaneous movement segments. Such segments in our complex actions gracefully build an ebb and flow of intended actions and sensory consequences [11]. Some of these sensations that voluntary movements give rise to [19], return to the brain as intentional feedback, thought to contribute to our internal models of action dynamics [20, 21]. This form of volitionally controlled kinesthetic reafference cumulatively helps us build accurate predictions of those intended sensory consequences [19], while other unintended movements return to the brain as spontaneous reafference, providing contextual cues that support motor learning, motor adaptation and action generalization across different situations [11].

One informative aspect of this ebb and flow of intent and spontaneity in our actions is the fundamental differences that emerge in the geometric features of the positional trajectories that the moving body describes [22–24]. When the motions are intended, geometric invariants derived from these trajectories emerge and remain robust to changes in speed dynamics [16, 23–27]. In contrast, trajectories from unintended motions produce different signatures of motor variability bound to return to the brain as spontaneous feedback. These internal sensations help us define contextual variations emerging from external environmental cues [11, 12]. These may include for example, changes in visual and auditory inputs, such as shifts in lighting conditions, or modulations in sound and music [12, 28]. The geometry of these spontaneous movements’ trajectories dramatically changes with fluctuations in the movements’ dynamics. Changes in speed [16, 23–27] or mass [12] affect their motor variability in fundamentally different ways (if we compare the signatures of variability derived from the spontaneous samples to those derived from deliberately staging the same movement trajectories [12, 25].) More importantly, the fluctuations in the motor variability of these spontaneous motions can forecast symptoms of Parkinson’s disease before the onset of high severity [29, 30]. They have also aided in evoking the sense of action ownership and agency in young pre-verbal children [31]. For these reasons, here we posit that deliberate and spontaneous segments of complex actions ought to differentially contribute to our sense of action ownership and to our overall sense of agency. To examine this proposition, we follow a phylogenetically orderly taxonomy of the nervous systems’ maturation (Figure 1B) and examine all levels of neuromotor control – from autonomic to deliberate – necessary to coordinate voluntary motions (Figure 1A).

More specifically, since autonomic systems are vital to our survival and wellbeing, they may remain impervious to subtle distinctions between deliberate and spontaneous motions that take place across the body, as the end effector completes goal-directed actions. Here we explore the interplay between autonomic signals and voluntary motor control in actions that integrate deliberate and spontaneous motions across the body. We use a new unifying statistical framework for individualized behavioral analyses and network connectivity analyses and offer a quantitative account of how these movement classes contribute to the overall embodied sense of agency.

## 2. Materials and Methods

### 2.1 Experimental Design

#### 2.1.1. Participants

Nine undergraduate students (2 males and 7 females) between the ages of 18 and 22 years were recruited from the Rutgers human subject pool system. Two were left-handed and seven were right-handed, and all had normal or corrected-to-normal vision. All participants received credit for their participation, and provided informed consent, which was approved by the Rutgers University Institutional Review Board. The study took place at the Sensory Motor Integration lab at Rutgers University.

During the experiment, movement kinematics and heart signals were recorded from each participant. However, one participant’s recording had too much noise (i.e., inaccurate sensor position with error larger than 10cm), so we excluded this participant’s data in the analysis. For that reason, eight participants’ motor and heart signals were analyzed.

#### 2.1.2 Sensor Devices

##### Motion capture system (kinematics data)

Fifteen electromagnetic sensors sampling at a frequency of 240 Hz (Polhemus Liberty, Colchester, VT) were attached to the participant’s upper body in the following locations: center of the forehead, thoracic vertebrate T7, right and left scapula, right and left upper arm, right and left forearm, non-dominant hand, and the dominant hand’s index finger. These sensors were secured with sports bands to allow unrestricted movement during the recordings. Motor signals were recorded in real-time by Motion Monitor (Innovative Sports Training Inc., Chicago, IL) software, where the participant’s body was constructed by a biomechanical model, and movement data were preprocessed by an embedded filtering algorithm of the software, providing the position and kinematics of each sensor.

##### Electrocardiogram (heart data)

Three sensors of electrocardiogram (ECG) from a wireless Nexus-10 device (Mind Media BV, The Netherlands) and Nexus 10 software Biotrace (Version 2015B) were used to record heart activity. At a sampling rate of 256Hz, the sensors were placed across the chest according to a standardized lead II method.

#### 2.1.3. Experimental procedure

Participants sat at a desk facing an iPad tablet (Apple, Cupertino, CA), which was used to display stimuli during the experiment, and participants responded by touching the tablet screen. The tablet display was controlled with an in-house developed MATLAB (Release 2015b, The MathWorks, Inc., Natick, Massachusetts, United States) program and TeamViewer application (Germany).

As shown in Figure 2, for each trial, the participant was presented with a circle on the tablet screen. This circle served as a prompt for the participant to touch the tablet screen within five seconds. After the touch, either 100ms, 400ms, or 700ms elapsed, and the participant heard a tone at 1000Hz for 100ms. Then, on the tablet screen, the participant was presented with a sliding scale, ranging from 0 to 1 (second), to indicate how long he/she perceived the time elapsed between the touch and the tone. The response was to be made within five seconds upon the display of the sliding scale. The five seconds time-window was considered enough for the participant to provide a response, as it took approximately 1 s to touch the screen and retract the hand back to its original position. There was a total of three conditions – control, low cognitive load, high cognitive load - and each condition consisted of 60 trials. In the control condition, the participant simply performed each trial with no additional task; under the low cognitive load condition, the participant performed each trial while repeatedly counting forward 1 through 5; under the high cognitive load condition, they counted backwards from 400 subtracting by 3 while they performed each trial. Participants counted forward and backward at their own comfortable pace, and they took breaks in between each condition. The experiment set up took about 30 minutes, and the recording took about 40 minutes.

**Figure 2.**
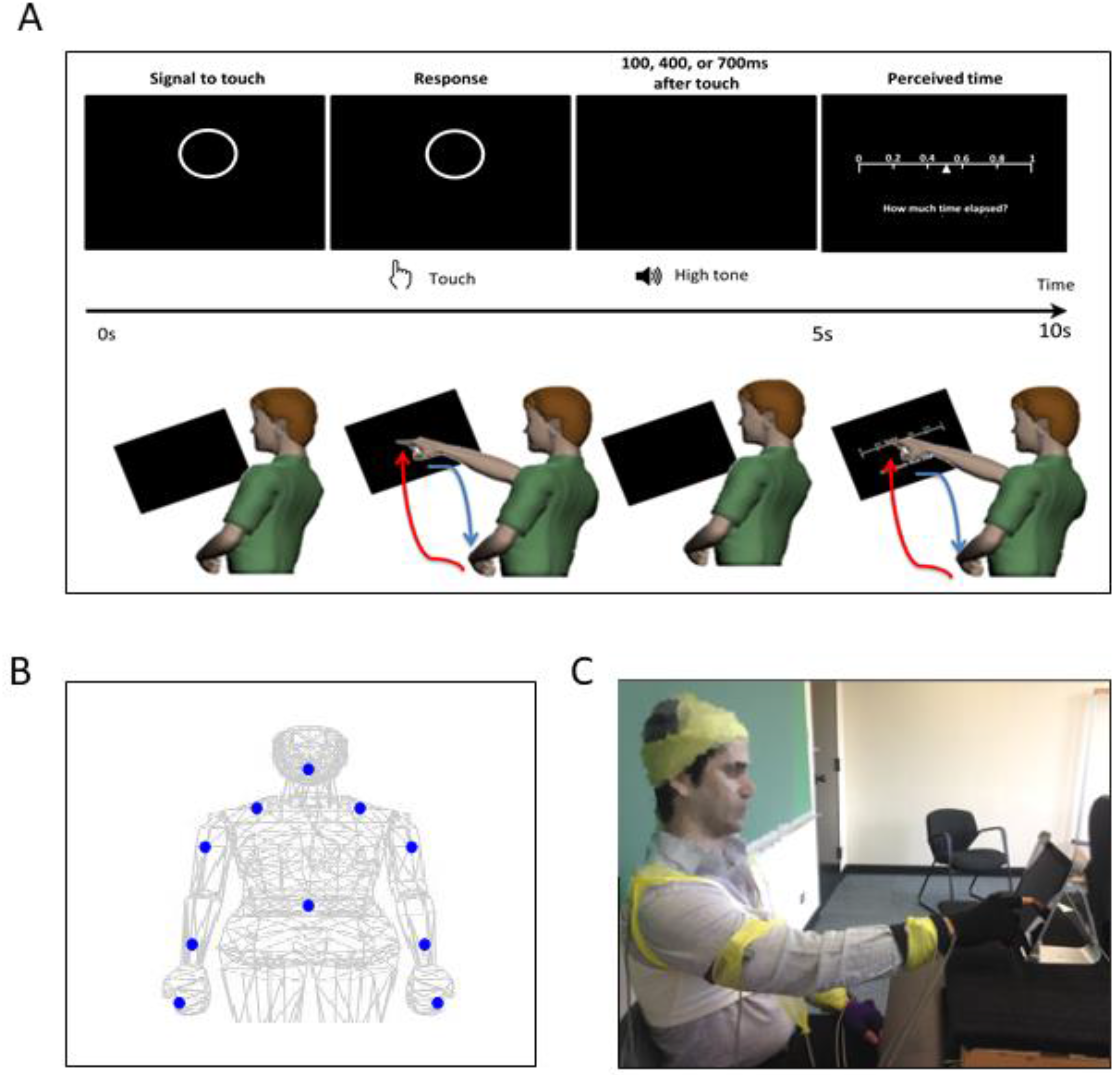
Experimental assay and instrumentation setup. **(A)** Experimental procedure. In a single trial, the participant was presented with a display screen as shown on the top panel. During the first 5 seconds, the participant was presented with a circle as a prompt to touch the circle on the screen. After the touch, the participant heard a tone. The duration between the touch and the tone was randomly set to be 100ms, 400ms, or 700ms. In the next 5 seconds, the participant was presented with a sliding scale, where s/he indicated how long the time was perceived to have elapsed between the touch and the tone, by touching the corresponding number on the scale. For each trial, the participant made two pointing gesture – one to touch the circle and another to indicate their time estimation on the sliding scale. Such pointing gesture was composed of a forward reaching segment (red) and a backward retracting segment (blue), as shown in the bottom panel. **(B)** Motion capture sensor positions. The sensors were attached on the following body parts: center of the forehead, thoracic vertebrate T7, right and left scapula, right and left upper arm, right and left forearm, non-dominant hand, and the dominant hand’s index finger. **(C)** Snapshot of the experiment. During the experiment, the participant was seated in front of the tablet screen to perform the tasks, and wired sensors were secured with athletic tape.

### 2.2. Statistical Analysis Overview

#### 2.2.1. Preprocessing

In this study, we extracted the kinematics (i.e., linear speed, angular acceleration) and heart data during time segments when the participant made a pointing motion towards the circle presented on the tablet screen; and combined them across the three conditions. As a result, we analyzed the kinematics and heart data recorded while the participant made 180 pointing motions (less any trials that were deemed noisy; the most trials we excluded per participant due to instrumentation noise were 12 trials).

To analyze the ECG and kinematics data in tandem, we up-sampled the kinematics data from 240Hz to 256Hz using piecewise cubic spline interpolation. Note, the ECG signals were not synchronized with the kinematics data but were manually time stamped at the start and end of each experimental condition. For that matter, we expect a presence of lag between the two modes of signals – kinematics and ECG – but the lag would not exceed 1 second.

To exclude effects of muscle motion from the ECG heart data, we bandpass filtered the data with Butterworth IIR for 5-30Hz at 2nd order. This filter was effective in identifying QRS complexes and extracting R-peaks in previous studies [13, 32]. Here, the filter excluded the dominant frequency range where typical kinematics signals are present (see Appendix Figure A1). We performed our analyses using both filtered and non-filtered EKG data and found similar trends and patterns. However, the paper only presents the results from using the filtered data, as it is a better reflection of the heart activity.

#### 2.2.2. Data analysis structure

We used the rationale in Figure 1 to structure our analyses, with a focus of two main axes denoting the level of motor intent and awareness that the brain may have during complex tasks (Figure 3A). More precisely, one axis explores possible differentiations between time segments of the pointing movements that are deliberately aimed at an external target (forward/high motor intent) vs. segments that are consequential to the deliberate ones (backward/low motor intent). The latter may occur when the hand retracts back to rest, or when after touching the target the person transitions the hand in route to another goal-directed motion. These segments have been studied in our lab across very complex motions in sports (boxing, tennis) and in the performing arts (ballet, salsa dancing). We have coined them spontaneous movements and discovered that they have precise signatures that distinguish them from the deliberate ones. For this reason, we hypothesized here that these spontaneous motions would have different stochastic signatures or be differentially expressed in relation to the deliberate ones.

**Figure 3.**
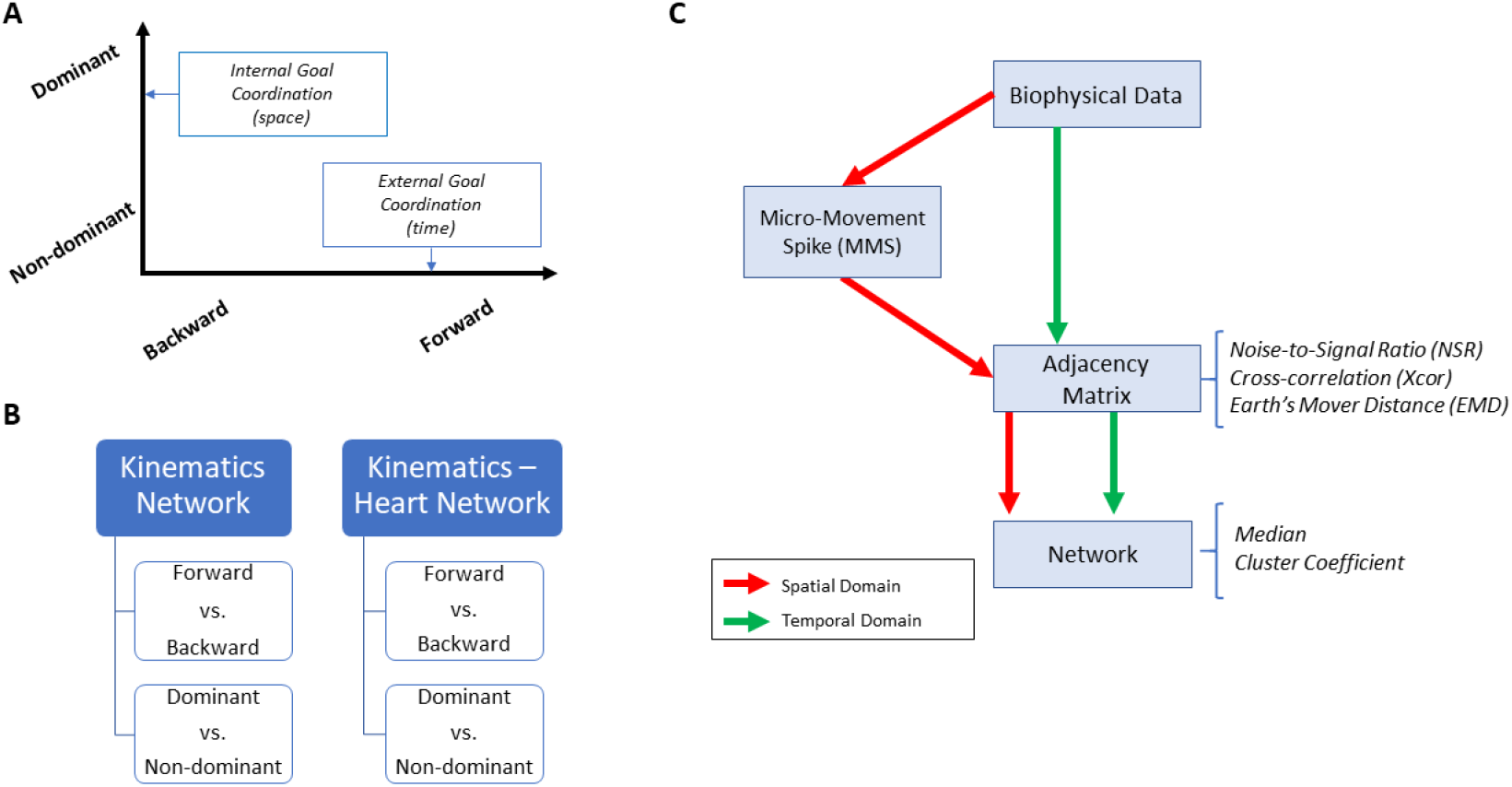
Overview of analytics pipeline. **(A)** Behavioral assay to quantify ranges of motor intent along two axes to highlight externally and internally defined goals. Along the former, motions are classified across time based on the end-effector’s movement, ranging from backward-spontaneous (lower motor intent) to forward-deliberate (higher motor intent) motions. Along the other axis, motions are classified across locations of the body, based on the proximity to the end-effector, from non-dominant side of the body parts (lower motor intent) to the dominant side including the end-effector (higher motor intent). Note, the two axes are not necessarily orthogonal as the schematics imply. **(B)** Two types of network analyses were made. Within the kinematics network, kinematics data served to compare patterns of variability from movement segments of higher level of intent (deliberately aimed at the goal) and movement segments with lower level of intent (spontaneous retractions of the hand to rest, without instructions), including as well comparison of patterns from the dominant and non-dominant parts of the body. Within the kinematics-heart network, a similar comparison was made, with a layer of autonomic function added, using signals from the EKG sensors. **(C)** For the spatial domain of connectivity analysis, raw biophysical data (biorhythms) co-registered from multiple layers of the peripheral and autonomic nervous systems were converted to MMS, and used to compute pairwise similarity/synchronicity metrics to build adjacency matrices to represent weighted / undirected graphs. For the temporal domain, the raw biophysical data were directly used to build adjacency matrices. For both domains, with the obtained adjacency matrices, network connectivity analyses combined with non-linear dynamical systems approaches were used to identify self-emerging kinematic synergies and various indexes to enable objective quantification of the embodied cognition phenomena.

The other axis explores possible contributions of body parts that are not directly related to the end effector (the dominant hand) performing the pointing task. We reasoned that there may be higher motor intent devoted to the performing (dominant) hand of the participant than to the non-dominant side of the body. Furthermore, we explored how other body parts (also co-registered within the sensors’ network) contributed to the overall performance of this task.

These two axes were explored at the voluntary level of motor control interleaving deliberate goal-directed (forward) actions and spontaneous (backward) segments of the full pointing loop. We also included in our analyses the autonomic level of control in the taxonomy of Figure 1A. And to that end, we co-registered the heart activity and incorporated it into the bodily kinematics activity (Figure 3B). We next explain how to overcome challenges in sensors’ data fusion from disparate systems along with new approaches to analyze these multi-modal data.

#### 2.2.3. Challenges of multilayered data with non-linear dynamics and non-normally distributed parameters

##### Disparate physical units

Different instruments to assess biorhythms from different layers of the nervous systems (i.e., kinematics *vs*. EKG) output biosignals with different physical units (e.g. *m/s* from the kinematics speed, *mV* from the EKG). This poses a challenge to integrate these signals and examine their interrelations across these layers.

##### Allometric effects

Another issue is that when examining such data from different participants with different anatomical sizes, allometric effects may confound our results. This is so because *e.g*. the speed ranges that a person attains depend on the length of the arm. Longer arms tend to broaden the ranges of speed and contribute to the distribution of speed values that the person attains in any given experiment. As such, we need to account for these possible allometric effects.

##### Assumption of normality

Another related matter to the ranges of speed and their distributions is that they vary from person to person according to multiple factors (*e.g*. age, body mass, sex, fitness, etc.) [32]. These variations result in probability distributions with heavy tails, which are incompatible with common assumptions of normality in the literature. When the effects of the task, or the inherent motor noise in the system, are such that most values related to the speed distribute more densely toward the left of the frequency histogram (*e.g*. in autism exponentially distributed maximum speed amplitude is common [33]), assuming normality may incur in spurious results. This is so, because speed ranges from 0 to some limiting value for each person (the maximum speed that the person can reach before damaging the joints). As such, when one obtains the mean +/- two standard deviation values to approximate standard error bars (which is very common in the motor control literature) while summarizing the statistical features of the data, the data may fall in the negative speed ranges (which is physically absurd).

##### Assessing similarity in probability space

Going beyond significant hypothesis testing models, one may need to assess the differences between probability distributions. To that end, one may need a proper similarity metric. Yet, when our data represents points in probability space, and the distributions are not symmetric, it is challenging to assess their similarity in a consistently proper way. Measures like the Fisher information metric are designed to compare symmetric distributions and the Kullback-Leibler divergence is computed asymmetrically between distributions (one-sided). We would like to have a proper (two-sided) distance metric to assess change and its rate when points are related to non-symmetric continuous probability density functions, or to their discrete approximations.

##### Degrees of freedom across intent levels of motor control

Multiple locations of the grid of sensors, co-registering biorhythms from different nervous systems, contribute differently to the overall behavior of the system. Some may be more directly related to action success, while others may provide support. Separating the bodily region within a kinematics-heart network can be challenging because of the non-linear dynamics of the interactive systems. Yet, most methods assume or impose local linearities to model such phenomena. Here we propose to approach this problem by treating the grid of sensors as a dynamically evolving weighted interconnected network, whereby we track self-emerging modules informing us of spontaneous synergies and connectivity patterns.

#### 2.2.4 Some solutions to the Challenges

##### New data type for disparate physical units

We have created a data type called the micro-movement spikes (MMS), which is a unitless, standardized waveform derived from the moment to moment fluctuations in the raw data peaks’ amplitude and / or timing. This data type extracts the fluctuations in amplitude and/or timing of any waveform with peaks and valleys (e.g. time series of speed values or kinematic related values derived from them). To that end, we obtain the empirically estimated moments from the peaks in the raw waveform. We then build a new waveform that can be normalized according to various criteria. This new waveform is then unitless and refers to a relative quantity (rather than to an absolute quantity).

##### Data standardization to account for allometric effects

The Anthropology and Paleontology literature has several solutions to address comparative data that may come from different bone sizes across *e.g*. different humanoids [34, 35]. Equation (1) provides an example of standardization to scale values derived from any waveform with peaks and valleys, which can be derived *e.g*. from data series with different physical units, from effectors of different sizes:

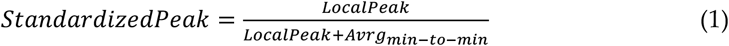

The standardized quantities are in the real-valued [0,1] interval. They are coined MMS amplitudes and treated as a continuous random process. We have characterized several complex behaviors from various layers of the nervous systems using the MMS, and expressed them in two forms: (1) without preserving the original frames of the data, *i.e*. just focusing on the MMS amplitude fluctuations and (2) conserving the original frames, in which case, we would 0-pad those that are not spikes, or preserve their values as additional gross data contributing to the phenomena in question. Either way, these fluctuations ought not be averaged out by assumptions of normality. Whereas in the extant literature these fluctuations are considered noise, or superfluous, here we treat them as important signal.

##### Distribution-free approach to counter current assumption of normality

We do not assume normality in the data. Instead, we gather enough data to empirically estimate the best family of probability distributions that fits the data. To that end, we here use maximum likelihood estimation (MLE) with 95% confidence intervals and seek the best continuous family that fits our data.

##### Distance metric to assess similarity in probability space

We here introduce the use of the Earth Mover’s Distance Metric (EMD) [36–39] to approximate (using the frequency histograms of the MMS amplitudes) the stochastic shifts in probability space that occur for different movement types. This is an appropriate similarity metric that allows us to examine the extent to which different levels of motor control change the stochastic patterns. We briefly describe it below:

The EMD, also known as the Kantarovich-Wasserstein distance [40], measures the distance between two discrete probability distributions. Given two discrete distributions P = {(p_1_,w_p1_), … (p_m_,w_pm_)} [13, 14], where pi is the cluster representative and wpi is the weight of the cluster; and Q = {(p_1_,w_p1_), … (p_m_,w_pm_)}, EMD computes how much mass is needed to transform one distribution into another. Defining D [d_ij_] as the ground distance matrix, where dij is the ground distance between clusters p_i_ and q_j_, and F = [f_ij_] with f_ij_ as the flow between p_i_ and q_j_; EMD is computed by minimizing the overall cost of such:

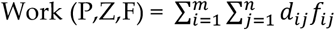

As there are infinite ways to do this, the following constraints are imposed to yield EMD values:

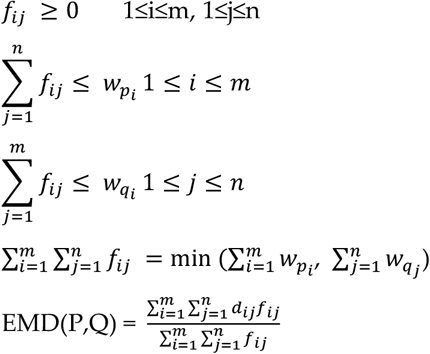

##### Network connectivity analyses to assess degrees of freedom recruitment across modalities of motor control

We use graph theory to examine the inter-relations across the nodes of the multilayered kinematics-heart network. To that end, we derive an adjacency metric of pairwise quantities reflecting the cross-correlation between any pair of nodes in the grid. We then construct weighted directed networks and borrow connectivity metrics from brain-related research. We extend these methods to represent the peripheral network using the bodily biorhythms from multiple layers of the nervous systems’ functioning, spanning from voluntary to autonomic (Figure 1A).

#### 2.2.5. Choice of kinematics parameter

The recording of positions over time across 10 upper body parts allows us to estimate two aspects of the biorhythmic data: *spatial* and *temporal* aspects, both of which are critical to characterize proper coordination and control. A parameter encompassing both aspects is the velocity. The derivative of position over time creates vector fields with direction and extent. Each point in the field (along the velocity trajectory) occurs in time and moves in space.

To assess spatial components, we use the scalar speed (distance traveled per unit time, where the unit time is taken constantly at the rate of 240 frames per second). We use Euclidean norm to compute the length of the velocity vector at each unit time, thus quantifying the rate of change in position per unit time – the linear speed (*m/s*). Likewise, we use the orientation data from each sensor and obtain the angular velocity from the rotations of each body part. Using appropriately the quaternion representation of rotations and the Euclidean metric to quantify the magnitude of the angular velocity vector, we obtain the angular speed (*deg/s*). These waveforms derived from the first order change are useful, but at the time scale (~1/2 hour) of our experimental assay, they provide fewer peaks per trial than waveforms derived from the second order change (*i.e*., linear acceleration (*m/s^2^*) or angular acceleration (*deg/s^2^*)).

As we need many spikes for our distribution-fitting and stochastic analyses, we used the angular acceleration kinematics data. Note, it is possible to have had participants perform more trials to obtain a larger number of spikes using the linear speed; however, this would fatigue the participants as the length of the experiment is around 70 minutes (inclusive of 40 minutes for set up). For that reason, within this amount of time, it was ideal to use the angular acceleration as our kinematic parameter of interest. This choice of parameter to analyze the stochastic patterns of the moment by moment fluctuations in signal amplitude (*i.e*. the spatial component of our analysis) provides tighter confidence interval in the empirical estimation of the best probability distribution family fitting the data.

We also examined temporal components of the data. To that end, we used the linear speed patterns and the cross-correlation function. We extended our analyses to different kinematics parameters, and while they all showed similar patterns and trends, we found the linear speed to best characterize the differing patterns of motor intent. For that reason, we present the results of the temporal analyses involving cross-correlation based network connectivity patterns using the linear speed as our waveform of choice (Figure 3C).

### 2.3. Data analysis on kinematics network connectivity

As a first step, we separated the kinematics data obtained from all 10 body parts, using the start and end time of the dominant hand making a forward-deliberate motion, and the hand making a backward-spontaneous motion (Figure 4A). This is possible to do (automatically) because (1) the speed is near 0 at the onset of the motion towards the target; (2) the distance to the target monotonically decreases and once again the hand pauses at the target at near 0 speed. As the deliberate (forward) segment is completed, the speed rises again away from 0 and the distance to the target increases as the hand follows the backward segment of the full pointing loop. The two segments can be automatically differentiated also because the deliberate (forward) one is less variable than the spontaneous (backward) one [11, 29, 34, 42].

**Figure 4.**
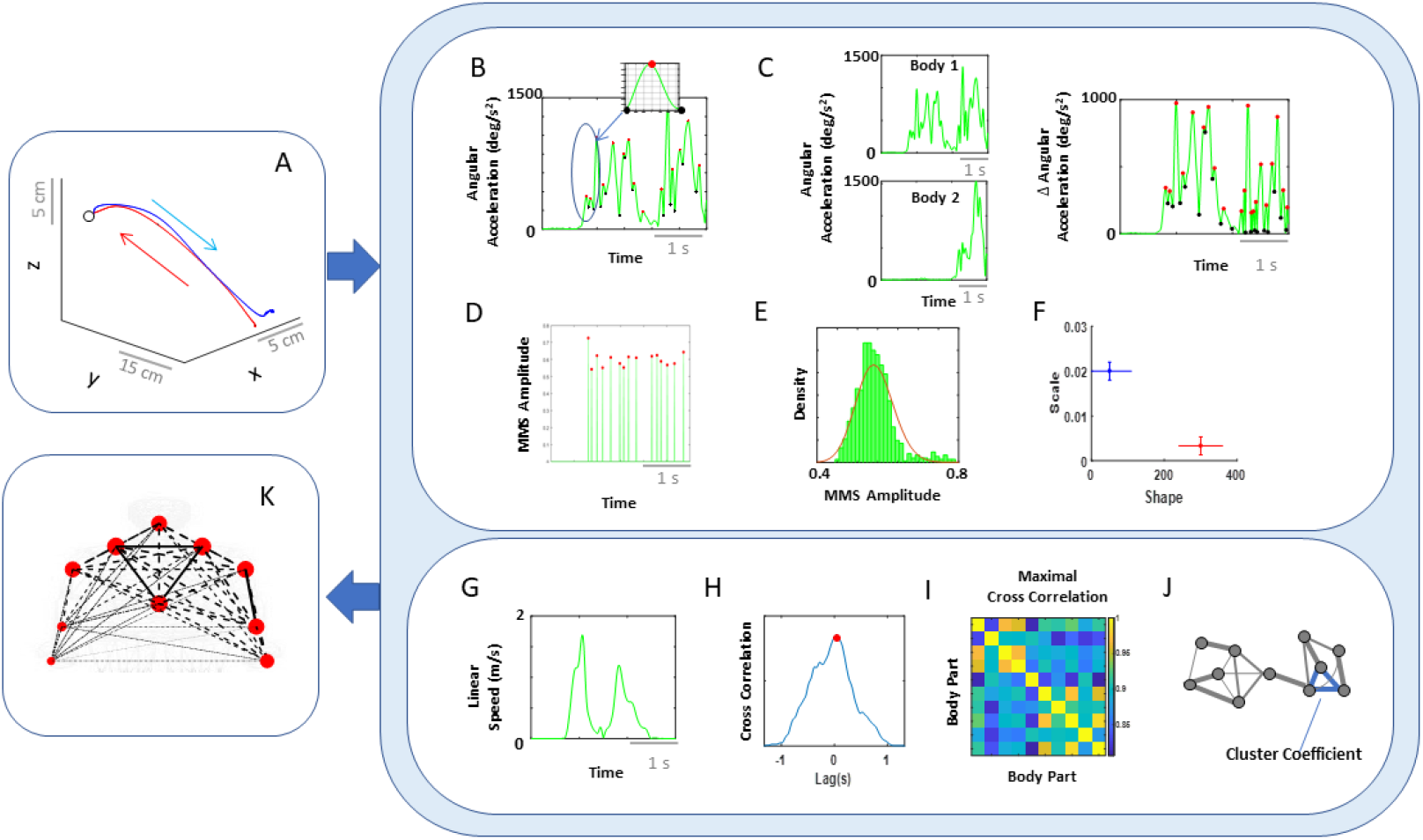
Analytical pipeline and visualization methods for the kinematics network. **(A)** Representative movement trajectory of the dominant hand during a pointing motion to a target (denoted by a small open circle). Each trial comprised of a forward-deliberate (red) and backward-spontaneous (blue) segment. These could be automatically separated by the speed and distance criteria (see Figure A2). **(B)** Time series of angular acceleration of the dominant hand’s index finger during a typical pointing task. To examine kinematics-based connectivity, we used the angular acceleration time series, focusing on the moment by moment fluctuations in waveform amplitude. Here, peaks (maxima) and valleys (minima) are shown in red and black dots, respectively. The inset shows a zoomed-in picture of a single angular acceleration segment (i.e., two local minima and a single peak in between, used for standardization described in Eq 1). **(C)** Pairwise absolute difference in waveform was obtained and standardized using Eq 1. The resulting waveform provided the input to obtain MMS. **(D)** MMS train scaling the waveform amplitude for a typical pointing task. All standardized spike amplitude values from (B) and (C) were maintained, while all non-spike values were set to 0. **(E)** Frequency histogram of MMS amplitudes fitted to a Gamma PDF using MLE. **(F)** The empirically estimated Gamma parameters (shape and scale) were obtained and plotted on a Gamma parameter plane, with marker lines representing the 95% confidence interval. Noise-to-signal ratio (NSR) (i.e., fitted Gamma scale parameter) were later used for comparison between motor segments and dominance side. **(G)** Representative time series of linear speed of the dominant hand’s index finger in one trial. **(H)** Pairwise cross-correlation between two body parts. **(I)** Adjacency matrix obtained from all pairwise maximal cross-correlation across all body parts under consideration, to represent a weighted undirected graph. **(J)** Connectivity metrics (e.g. clustering coefficient) were used to quantify patterns of temporal dynamics. **(K)** Network connectivity analyses to unveil self-emerging clusters, where nodes correspond to each body part. For the spatial domain, NSR derived from MMS amplitudes of angular accelerations were visualized as node size, and NSR derived from MMS amplitudes of pairwise absolute difference in angular acceleration as edge thickness. For the temporal domain, cluster coefficients were visualized as node size, and median cross-correlations as edge thickness.

For the connectivity analysis centered on spatial aspects of the signal amplitude, we pooled the angular acceleration data from each body part and extracted the MMS amplitudes (referred to as MMS from hereon). We then built frequency histograms of the MMS and explored several families of PDFs using MLE. The continuous family of Gamma PDFs yielded the best fit (Figure A2) and served to provide the noise to signal ratio (NSR; computed to equal the Gamma scale parameter) for each body part (Figure 4B, 4D-F). These were then visualized as node size in the schematics of the network in Figure 4K across different motor intent levels.

To characterize the connectivity of 2 body parts, we took the pairwise absolute difference between angular acceleration and based on the obtained absolute difference time series, computed the corresponding MMS. We then fitted the Gamma scale parameter (i.e., NSR) (Figure 4C, 4D-F), which were visualized as edges in the schematics of the network in Figure 4K. The intuition behind taking the absolute difference in angular acceleration time series from two body parts is that this reflects the change in positional distance between those two body parts, and thus represents the connectivity (physical distance) between those two. The NSR values were then compared between different movement segments (i.e., forward vs. backward) and different hand dominance (i.e., right vs. left arm/hand), to understand the noise level during different levels of motor intent. Note, for each type of motor segment (i.e., forward vs. backward), and for each dominance side (i.e., dominant vs. non-dominant), more than 2500 spike amplitude data were extracted. These spike amplitude data were then plotted on a frequency histogram using Freedman-Diaconis binning rule [43]. They were used for empirical estimation of the best PDF in an MLE sense. The results yielded the Gamma probability distribution function (PDF) (see Figure A2 B).

Connectivity analyses on temporal aspects of coordination involved the linear speed from each pair of body parts. We computed pairwise cross-correlations to derive an adjacency matrix that would represent a weighted undirected graph. Here, the ij-link’s weight is the maximum cross-correlation value between nodes i and j (that is, the corresponding two body parts). From these matrices, we computed clustering coefficients, which are measures that characterize the local connectivity (i.e., functional segregation). They would represent self-emerging kinematic synergies. Specifically, the degree of a node in the network (number of links at a node) between a set of nodes form triangles, and the fraction of triangle numbers formed around each node is known as the clustering coefficient (Figure 4G-J). This measure essentially reflects the proportion of the node’s neighbors (i.e., nodes that are one degree away from the node of interest) that are also neighbors of each other [44]. Here, we computed the average intensity (geometric mean) of all triangles associated with each node, where the triangles reflect the degree strength, and is computed as shown below (using an algorithm by [45]; Eq 2).

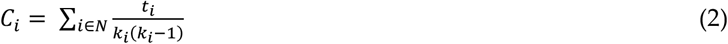

*N:* set of all nodes (composed of 10 body parts)

*C_i_*: cluster coefficient for node *i* (*i ∈ N*)

*t_i_*: geometric mean of triangles links formed around node *i* (*i ∈ N*)

*k_i_*: number of degrees (links) formed around node *i* (*i ∈ N*)

To visualize the network, we represented the median pair-wise cross-correlation values as the edge thickness, and median cluster coefficient values as the node size (Figure 4K). The median cross-correlation and cluster coefficient values were then compared between different movement segments (i.e., forward vs. backward) and different hand dominance (i.e., right vs. left arm/hand), to understand how linear correlations differed across varying levels of motor control.

### 2.4. Data analysis on kinematics-heart network connectivity

As with the kinematics connectivity analysis, we segmented the data of the filtered EKG data along with the kinematics data by the time intervals when the dominant hand was making a deliberate forward motion and a spontaneous backward motion. (Figure 5A).

For the spatial domain of connectivity, we took the segmented data of angular acceleration and EKG data, and extracted MMS from both signals, and plotted a histogram of the MMS. Because the MMS of EKG signals did not follow a Gamma distribution, in order to assess the connectivity between the two, we computed the earth mover’s distance (EMD) between the histogram from a single body part and from the EKG data (Figure 5A-D).

**Figure 5.**
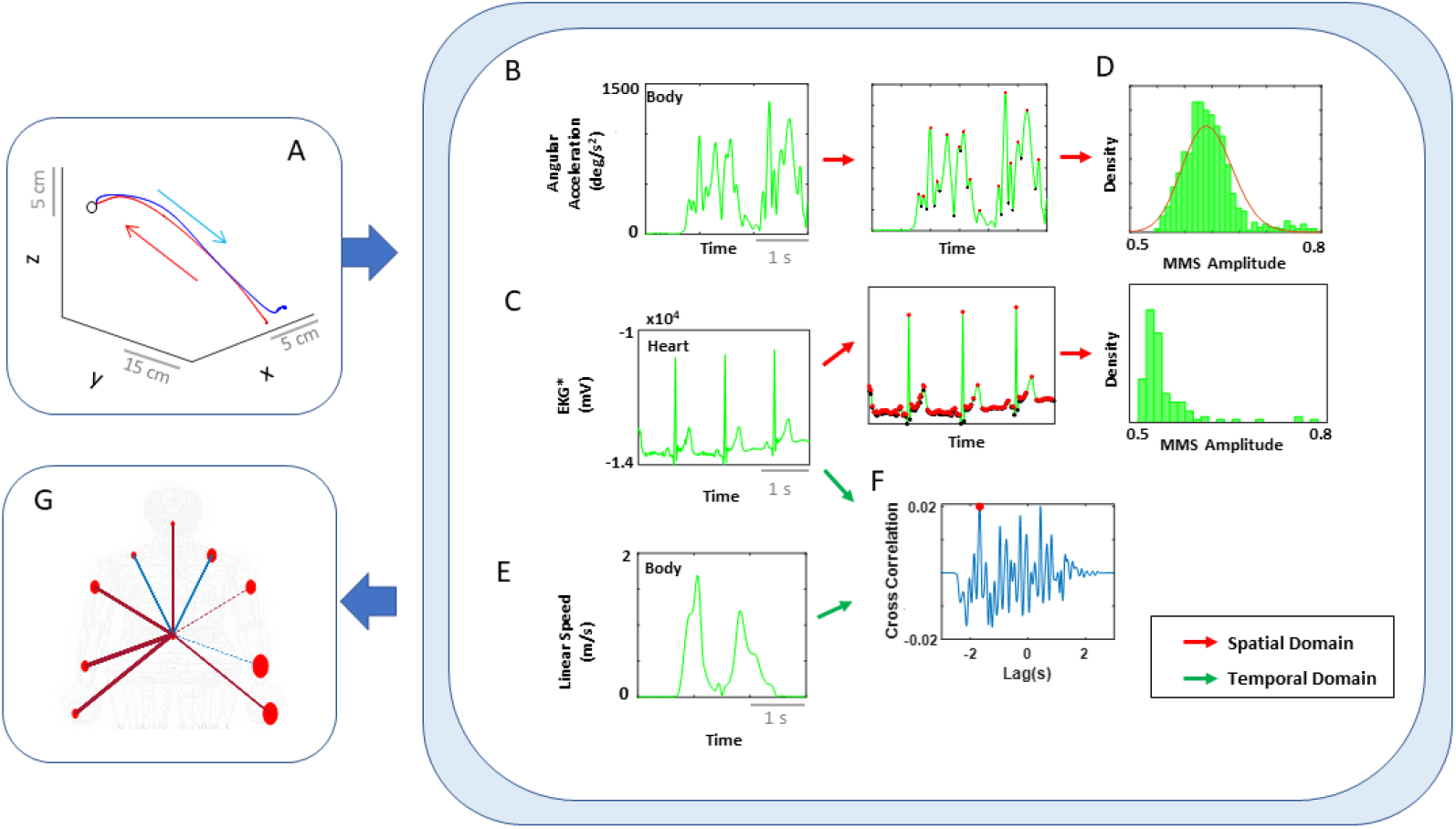
Analytical pipeline and visualization methods for the kinematics-heart network. **(A)** Typical movement trajectory of the dominant hand position, while performing a single pointing action towards a target. Each trajectory was separated into forward-deliberate (red) and backward-spontaneous segments (blue) according to hand-target updated distance and near-zero-speed value (see Figure A2 for details). **(B)** Angular acceleration time series of the hand during a typical pointing task. MMS amplitudes from the angular acceleration time series were extracted for each body part. **(C)** Filtered EKG time series during a pointing task. MMS amplitudes from the filtered EKG time series were extracted. **(D)** Histograms of compiled MMS amplitudes. For spatial analysis, pairwise EMD was computed between histograms from each body part and heart activity. **(E)** Linear speed time series of the dominant hand. For temporal analysis, linear speed kinematics time series was used. **(F)** Cross-correlation between a single body part’s linear speed and filtered EKG signal. For each trial, cross-correlation was computed between a pair of filtered EKG and a single body part’s linear speed time series, and the maximal value (red dot) and its corresponding lag values were extracted. **(G)** Visualization of connectivity. Network connectivity was visualized, where node size represented the EMD between the corresponding pair of body part and heart signals (i.e., spatial metric), and edge thickness represented the median cross-correlation values between the signal pairs (i.e., temporal metric). The edge colors were visualized, such that red would indicate EKG signals temporally leading linear speed signals, and blue would indicate linear speed leading EKG signals.

For the temporal domain, we computed pairwise cross-correlations along with its lag, between the EKG filtered time series and each body part’s linear velocity time series. In fact, in our analysis, we found an interesting pattern in directionality (i.e., lag) of correlation, and deemed informative to present them in the network graph. For that reason, edge thickness was represented by the median cross-correlation values, and color of the edges were visualized, where red would indicate EKG signals leading linear velocity signals, and blue would indicate linear velocity leading EKG signals (Figure 5G).

For all these metrics, we compared the medians between different movement segments (i.e., forward vs. backward) and different hand dominance (i.e., right vs. left arm/hand), to understand how stochasticity and temporal dynamics changed across varying levels of motor intent between the heart (from ANS) and kinematics (from PNS/CNS).

## 3. Results

### 3.1. Higher Motor Intent Results in Higher NSR in Spatial Parameters

Motor intent in the context of our experimental assay specifically refers to the level of deliberateness (or spontaneity) of the movement segment in route to an external target (away from it). An instructed pointing action to touch the target is a goal-directed reach with high level of intent. In contrast, the uninstructed spontaneous retraction away from the target carries lower motor intent than the goal-directed one.

As a first set of analysis, the MMS extracted from the angular acceleration data from each body part were aggregated across all trials and conditions, and arranged by different movement segments (forward-deliberate vs. backward-spontaneous) and different dominance side. The same was also done on the MMS extracted from the absolute difference in angular acceleration from all pairs of body parts. The NSR was found to be significantly higher when the motions were deliberate and on the dominant side. (Figure 6).

**Figure 6.**
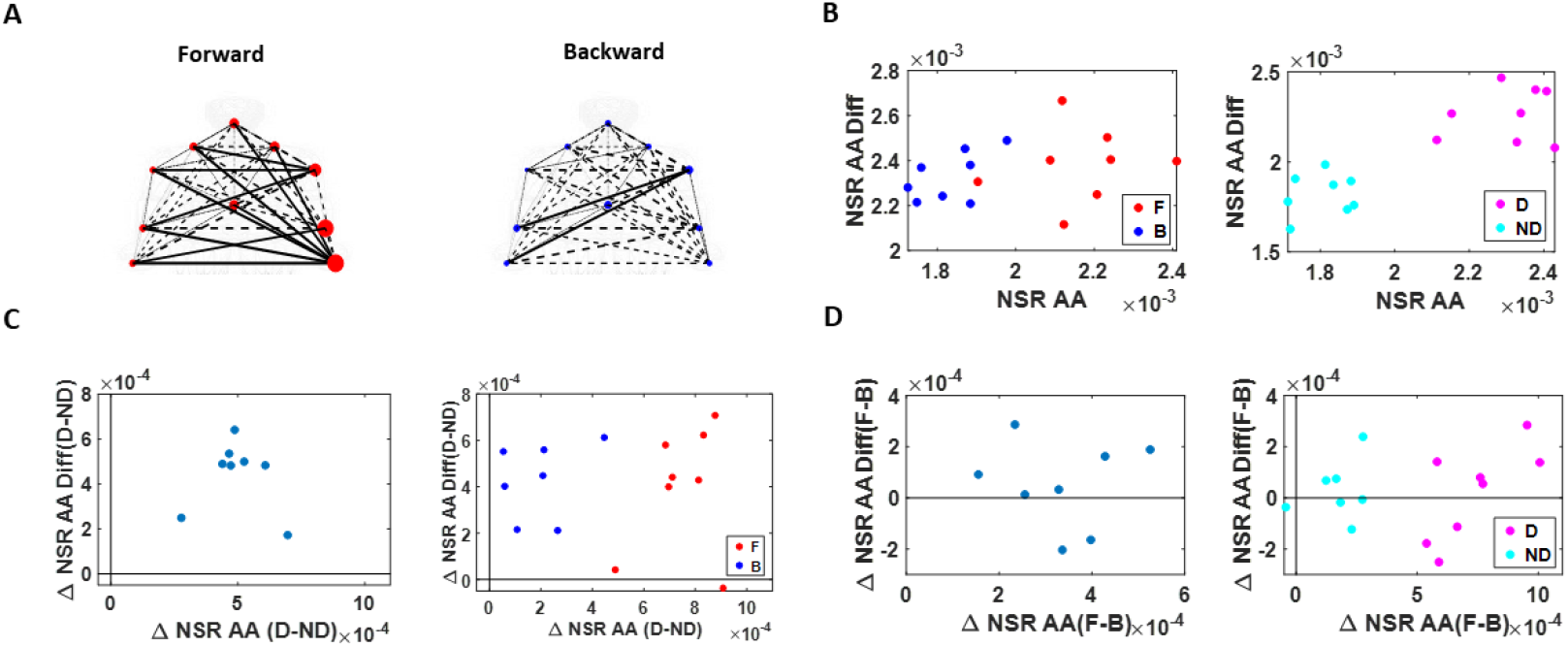
NSR signatures during pointing can differentiate the levels of intent. Comparison includes forward-deliberate *vs*. backward-spontaneous segments and dominant *vs*. non-dominant effector. **(A)** Network visualization of a right-handed representative participant. Node size is represented by the NSR derived from the corresponding body part’s kinematics time series, and edge thickness is represented by the NSR of the absolute difference in kinematics between the corresponding pairs of body parts. Node size and edge thickness are graphed in the same scale across different movement segments (i.e., forward and backward segments). **(B)** NSR for different movement segment and dominance side. Each dot is the median NSR values for each participant’s different movement segments (left) and dominance side (right) from the unitless MMS derived from the Angular Acceleration (AA) fluctuations in amplitude. The x-axis denotes the NSR from individual body part’s kinematics (NSR AA) and y-axis denotes the NSR from the MMS derived from the absolute pairwise body parts’ difference (NSR AA Diff). Generally, for the former (NSR AA) measure, NSR is higher during a forward segment (F; red) than during a backward segment (B; blue), and on the dominant side (D; pink) than on the non-dominant side (ND; cyan). **(C)** NSR difference between dominant *vs*. non-dominant side. Left panel shows the NSR median difference between the dominant and non-dominant side for each participant, denoted as a single marker. Right panel shows the NSR median difference between the dominant and non-dominant side for the forward motion (F; red) and backward motion (B; blue). When the difference between the dominant and non-dominant side is examined separately for each motion segment, the NSR AA difference is wider during forward motion segments (F; red) than during backward motion segments (B; blue). **(D)** NSR difference between forward *vs*. backward movement segment. Left panel shows the NSR median difference between the forward and backward motion segments for each participant, denoted as a single marker. Color scheme as in (B).

Specifically, NSRs of the kinematics time series from each body part showed was highest when an individual exerted higher motor control under higher level of motor intent, such as on the dominant side of the body and during a forward-deliberate motion. Conversely, when an individual did not deliberately intend to move the arm, as exhibited on the non-dominant side and during a backward-spontaneous motion, the NSR was at its lowest. The NSRs for all pairs of body parts’ absolute difference in angular acceleration (i.e., change in distance between the pairs of body parts), on the other hand, is higher on the dominant side (vs. non-dominant side), but does not show such consistent pattern when comparing between the two motion segments (forward vs. backward). Details of the 95% confidence interval of the fitted Gamma scale parameter (i.e., the NSR) for all participants, and for all body parts (Figure A3) and all pairs of body parts (Figure A4) can be found in the Appendix.

### 3.2. Higher Motor Intent Results in Higher Cross-correlations and Clustering of Temporal Parameters

We used the MATLAB Network Connectivity toolbox [26] and examined the adjacency matrix derived from the pairwise maximal cross-correlation coefficient based on the time series of linear speed values. The clustering coefficient (CC) was obtained for each body part as a metric of functional segregation. For analysis, we examined the median cross-correlation values as a function of the CC values. Here we found that higher level of motor intent (i.e. during forward-deliberate motion performed with the dominant hand) resulted in a tendency of increased CC and increased median cross-correlation values (Figure 7).

When we compared between different motion segments, median cross-correlations were higher for forward motions than for backward ones for all but two participants. When we compared between different dominance side, all participants showed higher correlation on the dominant side than the non-dominant one. The median CC showed to be higher for forward motions than for backward segments for all participants, and higher for the dominant side than the non-dominant side for all but two participants. For all participants, both measures showed statistical significance in their difference (see Table A1 of Appendix for detailed statistical results).

**Figure 7.**
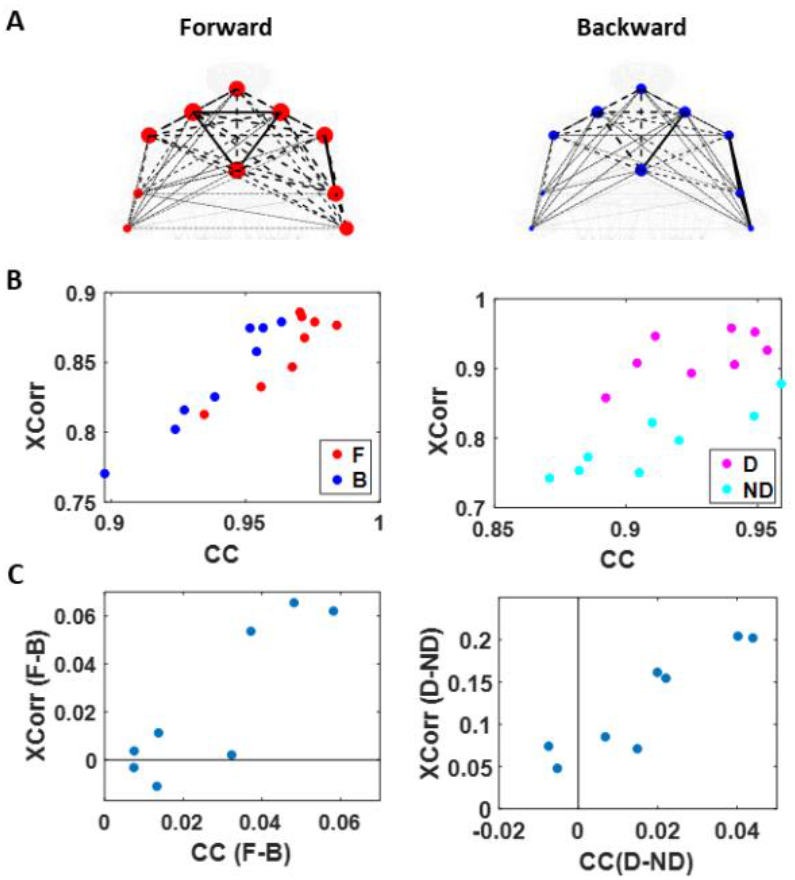
Network connectivity metric (cluster coefficient) and median cross-correlation differentiates between levels of intent. **(A)** Network visualization of a representative right-handed participant. Cross-correlation is represented by the line weight and cluster coefficient (CC) by the node size, during forward (left) and backward movement segment (right). **(B)** Median cross-correlation (y-axis; Xcorr) and CC (x-axis) of linear speed for each participant’s movement segment (left) and dominance side (right). Forward motions (red) and dominant side (pink) exhibits higher cross-correlation and CC values, than backward segments (blue) and non-dominant side (cyan). **(C)** Median cross-correlation and CC difference for different movement segments (left) and dominance side (right). Each participant’s data is denoted as a single marker. Higher motor intent tends to show higher cross-correlation and CC values.

The distinctions that we observe from these findings, on how different levels of motor intent have separable network connectivity patterns based on temporal aspects of the kinematics data, are consistent with the patterns uncovered using spatial aspects of the kinematics data. Specifically, when we exert higher intent on our body, regardless of the physical trajectory of the motion, there is a stronger connectivity across our body parts. However, we note that this pattern is not as uniform across all participants, as we had found in the spatial aspect of the network analysis.

### 3.3. Kinematics and EKG (heart) Signals Show Larger Stochastic Differences for Higher Motor Intent and Control

To assess patterns of connectivity between biophysical signals derived from voluntary and autonomic levels of motor control we examined the kinematics (generated by the CNS-PNS) and the heart activity (generated by the ANS). The patterns of MMS stochasticity and temporal correlation across these systems distinguished levels of motor intent and control.

The analyses involving EKG and kinematics revealed larger stochastic differences in MMS data when higher motor intent and control are exerted. More precisely, the pairwise EMD showed higher differentiation between these two signals in all but one participant when forward motion was made, but only on the dominant side of the body. Furthermore, all but two participants showed higher EMD on the dominant side of the body, but only during forward motions. On the other hand, however, when backward motion is made, we find an opposite pattern, where all participants show higher EMD on the non-dominant side. We infer that there may be a modulating factor that underlies the stochastic relation between kinematics and heart signals.

When we examine the temporal relations between the two signals, by computing pair-wise cross-correlations, we see higher cross-correlations when there is lower motor intent across all participants – that is, during backward motions, and on the non-dominant side. Here we note the low range of the correlation coefficient values, around 0.1. However, we see a similar trend when this is based on the non-filtered raw EKG data, with a higher range around 0.6.

### 3.4. EKG Leads Kinematics Under Higher Motor Intent, But Opposite Pattern Emerges in Spontaneous Motions Requiring Less Motor Intent

We also examined the lag values to assess which signal leads the other. We found that motions under higher motor intent (i.e., during forward-deliberate motions performed with the dominant side of the arm), EKG signals tend to lead the kinematics signal. On the other hand, in movements performed under lower intent (i.e., during backward-spontaneous motion, and on the non-dominant side of the arm), kinematics signals tend to lead the EKG signals. This is depicted in Figure 8.

**Figure 8.**
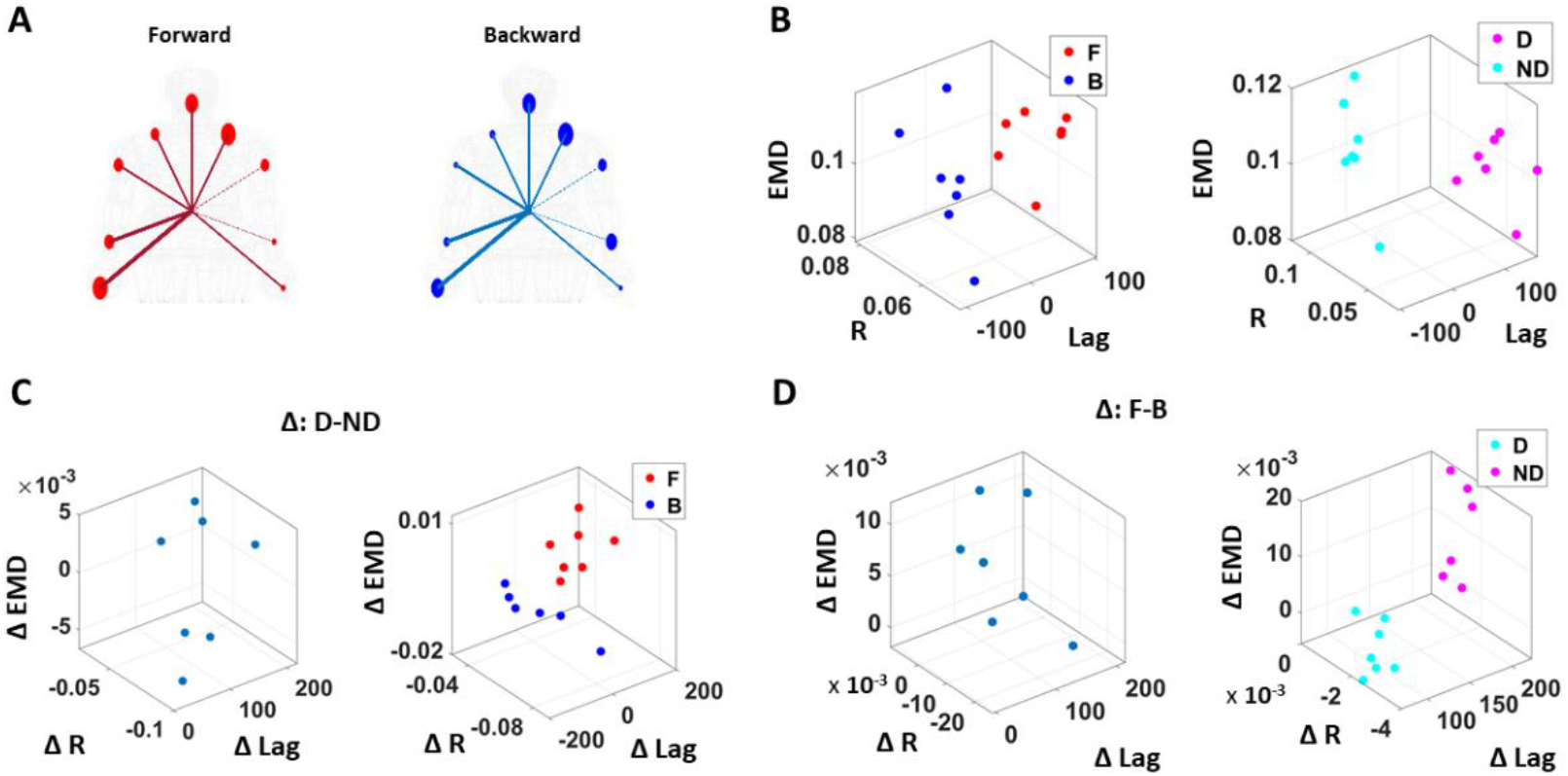
Differentiation of spatial and temporal connectivity within the kinematics-heart network according to levels of motor intent. **(A)** Network visualization of a right-handed representative participant. 1/EMD is represented by the node size, and median correlation is represented by the line weight. The color of edges indicates the temporal directionality between signals, where red indicates that heart leads the body linear speed and blue indicates that body linear speed leads the heart signals. **(B)** Median EMD (z-axis) and correlation (x-axis) and lag (y-axis) for each participant’s movement segment (left) and dominance side (right). There is an overall pattern where higher motor intent (denoted by red for forward motions, and pink for dominant side) is exhibited by lower correlations and EKG leading the kinematics signal (i.e., lag is positive value). **(C)** Median EMD, correlation, and lag difference for different dominance side (left), and this difference separated by movement segment (right). We find a pattern where pairwise EMD show higher differentiation under higher motor intent on the dominant side, but only when forward motion was made. **(D)** Median EMD, correlation, and lag difference for different movement segments (left), and this difference separated by dominance side (right). We find a pattern where pairwise EMD show higher differentiation under lower motor intent on the non-dominant side, but only when the backward motion was made.

We caveat that that because the EKG device and motion capture system was not exactly synchronized, the absolute lag value may not be as meaningful. Nevertheless, as we analyze these data in terms of the difference (i.e., the delta lag values between forward and backward motions, and between dominant and non-dominant sides), it is indeed meaningful to find such patterns uniformly across all participants.

Table1 summarizes the results that we showed in the sections above. We emphasize that although we examined a small number of 8 participants, each individual’s data is composed of a significant amount of data points with unique non-Gaussian stochastic characteristics. For that reason, instead of presenting the results with NHST (null hypothesis significant tests), we presented the results by comparing the median difference between data points from different levels of intent, for each individual.

**Table 1.** Summary of the connectivity results, where symbols^1^ are shown to indicate which category shows higher values.

|  |  | Kinematics (AA) Network |  |  |  |
| --- | --- | --- | --- | --- | --- |
|  |  | Forward | Backward | Dominant | Non-Dominant |
| Spatial | NSR AA | o |  | o |  |
|  | NSR AA Diff |  |  | o |  |
| Temporal | Cross-Correlation | $\Delta$ | | o | |
| | Cluster Coefficient | o | | $\Delta$ | |
|  |  | Kinematics (LS)-Heart Network |  |  |  |
|  |  | Forward | Backward | Dominant | Non-Dominant |
| Spatial | EMD | $\Delta$ (D)<sup>2</sup> | - | $\Delta$ (F)<sup>3</sup> | o (B)<sup>4</sup> |
| Temporal | Cross-Correlation | | $\Delta$ | | o |
|  | Lead*<sup>5</sup> | EKG | LS | EKG | LS |

<sup>1</sup> o indicates that it is higher for every participant; $\Delta$ indicates that it is higher for most participants
<sup>2</sup> Forward-deliberate motions have higher EMD only on the dominant (D) side
<sup>3</sup> Dominant side has higher EMD only during forward-deliberate (F) motions
<sup>4</sup> Non-dominant side has higher EMD only during backward-spontaneous (B) motions
<sup>5</sup> Lead\* shows which signal leads between the 2 signals

## 4. Discussion

This paper examined elements of the construct of agency from the embodied cognition framework and dissected several layers of neuromotor control contributing to the sense of action ownership. These layers, defined along a phylogenetically orderly taxonomy of maturation, follow a higher-to-lower gradient of intent, from voluntary, to involuntary, to autonomic signals. At the voluntary level, we followed the deliberate and the spontaneous segments of the target-directed pointing act, positing that they could differentiate between levels of intent and as such, delineate from the fluctuations in their biorhythmic activity, when a given movement segment was deliberately performed with intent vs. when the segment happened spontaneously without instruction. This differentiation is important to distinguish the sensory consequences of voluntary acts from those of acts that are not intended, or that occur autonomically. The sensory consequences of the latter are not currently studied, yet they seem important to complement von Holst’s and Mittelstaedt’s principle of reafference -as we know it today [19].

Our initial thought was that autonomic systems contributing to our brain’s autonomy over the body and to our overall embodied sense of agency would remain impervious to stochastic shifts at the voluntary levels. We reasoned that given the vital role of these systems for survival, their robust signal would not reflect subtle changes in levels of intent, motor awareness and voluntary control. As such, our guess was that if during voluntary movements, there were stochastic differences between deliberate and spontaneous segments of the reach, or between dominant and non-dominant sides of the body, such shifts in patterns of variability would not be appreciable in the heart signals’ fluctuations. Our guess was altogether wrong. Not only were the heart signals’ differences quantifiable at the level of fluctuations in signal amplitude; these differences were appreciable as well in the inter-dynamics of the kinematics and cardiac signals.

We found that when movements are intended and deliberately performed to attain the goal defined by an external (visual) target, the heart signal leads the movement kinematics signal. Yet, when these overt movements are spontaneous in nature, i.e. uninstructed and not pursuing the completion of a specific externally defined task goal, the heart signal lags the movement kinematics signal. Across spatial and temporal parameters, we found consistent trends and confirmed the trends through different parameters. Indeed, deliberate motions, performed with the dominant effector, carry higher levels of NSR, denoting higher fluctuations away from the empirically estimated mean.

We interpret these findings considering the principle of reafference [46]. Furthermore, we discuss the possible contributions of these self-generated signals to the self-emergence of cognitive agency from motor agency, namely, the sense that one can physically realize what one mentally intends to do, confirm the consequences (both intended and unintended) and as such own the action.

Von Holst and Mittelstaedt studied the complexities of reafference across the nervous systems in the 1950s. They tried to capture the inherent recursiveness that relates movements and their sensations as they flow within closed feedback loops between the external and the internal environments of the organism. They wrote, “Voluntary movements show themselves to be dependent on the returning stream of afference which they themselves cause.” And undeniably, feedback from voluntary movements currently play an important role in theoretical motor control, particularly within the framework of internal models for action [21, 47, 48] and more recent models of stochastic feedback control [49, 50]. Central to all these conceptualizations of the control problem has been the notion of anticipating the sensory consequences of impending intended actions. Nevertheless, nothing has been said about the consequences of action segments that bear a lower level of intent, that occur spontaneously, or that are altogether occurring autonomously. Modelers and experimenters in motor control do not seem to be aware of the former (although see [11, 12, 51]) and the latter are assumed to be far removed from cognitive processes (although see [13] more recently.) Yet, unintended consequences from the spontaneous segments of the voluntary action seem as important as those sensory consequences that result from the deliberate segments. They may serve to inform learning new tasks, adapting to new environmental conditions or situations and more generally, they may play a role as a surprise factor to aid propel curiosity and / or to stimulate creative, exploratory thinking. They may help make our “invisible” automatic movements visible to the conscious brain performing them, and/or to the external observer tracking our behaviors.

Neither these models, nor Von Holst’s work considered the contributions of unintended consequences from spontaneous acts quantifiable at different anatomical and physiological layers of the nervous systems, while trying to model the basic problem that the organism faces, i.e. the paradox of understanding the “self”, which entails parsing out external from internal reafference [52]. Without a unifying framework to quantify these multilayered interactions and their contributions to the emergence of the notion of self, it becomes rather challenging to bridge the cognitive sense of agency, and more basically of action ownership, “I can do this!; It’s me who’s doing this!” with the type of autonomous motor control that enables successful completion of the intended act. We argue that inclusion of the unintended consequences from overt spontaneous motions and autonomic signals in our models of motor control will help define embodied agency and provide a new framework to objectively quantify it.

The present work provides empirical evidence that (1) different levels of cognitive intent, awareness and control are indeed embodied and quantifiable in natural, unconstrained movements and (2) there are important contributions to central cognitive control quantifiable at the periphery in spontaneous segments of our motions and their consequences, but also in motions from supporting (non-dominant) body parts. Importantly, such differentiating contributions are also present in patterns from signals generated by the autonomic nervous systems. These aspects of the motor control problem are not considered at present in any of the mathematical and computational frameworks used to model the human brain, despite a body of empirical data differentiating classes of movements that are less sensitive to changes in dynamics [11, 16, 22–27] from those which are dynamic dependent [11, 12].

Our work augments Von Holst’s and Mittelstaedt’s principle of reafference nontrivially by including reafferent contributions from other layers of the nervous systems (Figure 1A) and highlighting the need to update our conceptualization of internal models for action [52]. In the past, the literature has focused on voluntary control and goal-directed behavior to define and to characterize agency [4, 9, 10, 53]. However, if new generations of AI models aim to attain artificial autonomous agents with real agency, it may be necessary to reformulate our models and reconceptualize our experiments in embodied cognition to encompass these multiple layers of intent, awareness and motor function.

An area of importance in this regard is smart health and AI, connecting digital biomarkers with clinical observational criteria (e.g. [54].) In the clinical world, there are many problems that will require to be mindful of this intended vs. unintended dichotomy, as there are phenomena that occurs spontaneously and is difficult to model within the voluntary reafference framework. The type of reafference that we need to model those problems belongs in the realm of self-emerging behaviors. Among these are sudden freezing of gait in Parkinson’s disease, leading to the loss of balance and occasional falls; seizures across a broad range of disorders; heart attacks; a subset of repetitive behaviors and self-injurious or aggressive episodes in autism, among others. All these episodes have in common the element of surprise connected to their spontaneity. No algorithm relying exclusively on intentional control signals can appropriately capture the essence of these phenomena. To properly characterize it, forecast it and quickly detect it, we need veridical generative models that understand the differences between the consequences of something that was intended and under voluntary control, something that spontaneously happened, and something that happens autonomically, with high accuracy. We do not have autonomous robots with embodied agency yet, because their staged motions are mostly pre-programmed. These programs may only mimic the predictive consequences of voluntary actions. Self-correcting robotic systems where such behaviors spontaneously self-emerge, are less common. It is perhaps self-emerging awareness derived from the consequences of spontaneous and autonomic phenomena that makes our embodied agency a special human trait contributing to intelligent control. This type of control combining deliberate and spontaneous acts, may produce solutions that are capable of generalizing from a small set of specific situations; transfer the learning from one context to another (using contextual variations) and retain robustness to potential interference from new situations in unknown contexts. In future research, it will be important to understand how the type of differentiation that we discovered here, paired with externally vs. internally generated rewards, may contribute to the fast or slow acquisition of memories from transient acts vs. memories from systematic periodic repetitions of those acts.

Here we offer a unifying framework with a taxonomy of function and differentiable levels of intent, awareness and control paired with a new statistical platform for personalized analyses of natural behaviors. This new model aims to capture and characterize the micro-fluctuations in the gross data of our biorhythms that traditional approaches throw away as noise through grand averaging and “one size fits all” methods. Our approach allows integration of multilayered hierarchical signals and provides the means to differentiate re-entrant contributions from multilayered exo- and endo-afference. This can help our self-realization of embodied agency as the spontaneous transformation of mental intent into physical volition. We invite the reader to consider this new model for embodied cognition and offer novel avenues to bridge the currently disconnected fields of motor control and cognitive phenomena.

## Author Contributions

Conceptualization, J.R. and E.B.T.; methodology, J.R. and E.B.T.; formal analysis, J.R. and E.B.T.; investigation, J.R. and E.B.T.; writing—original draft preparation, J.R. and E.B.T.; writing—review and editing, J.R. and E.B.T.; visualization, J.R. and E.B.T.; supervision, E.B.T.; funding acquisition, E.B.T. All authors have read and agreed to the published version of the manuscript.

## Appendix A

**Figure A1.**
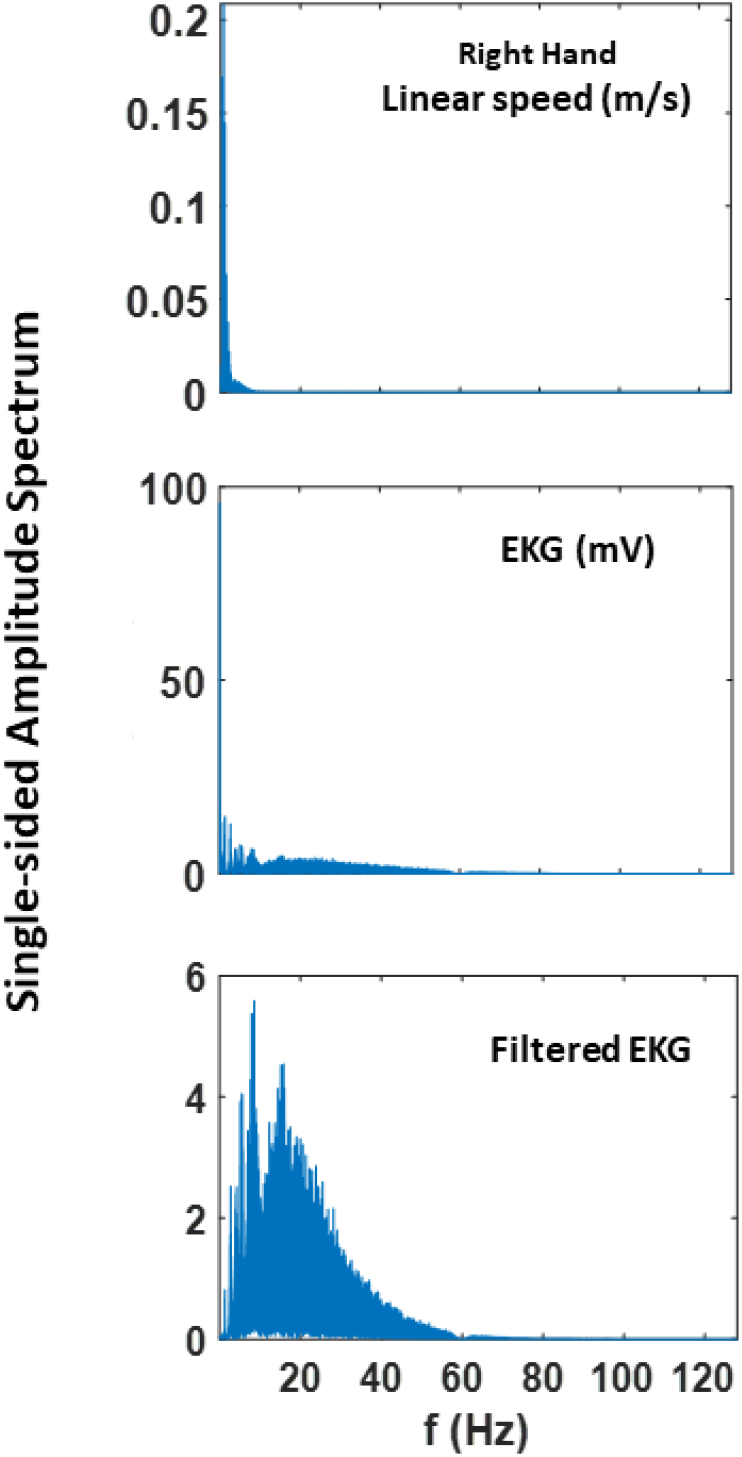
Fourier power spectrum of linear speed and EKG and filtered EKG signals extracted from 60 trials of pointing motion (i.e., 300 seconds).

**Figure A2.**
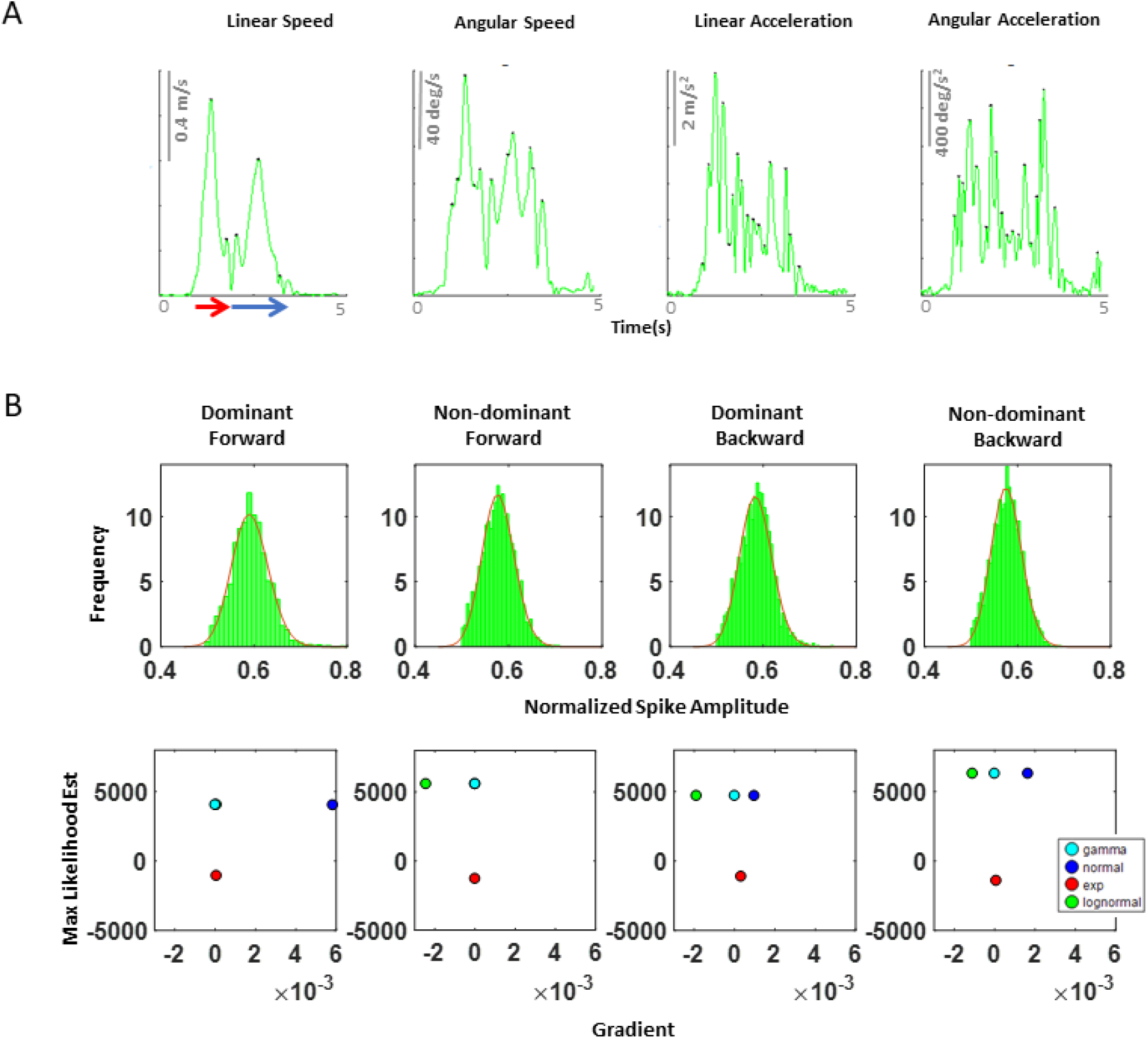
Speed profile of a typical pointing motion. During a single pointing motion, a typical speed profile of linear speed, angular speed, linear acceleration, and angular acceleration are exhibited as such. Because angular acceleration shows to have the largest number of peaks during a single pointing motion, we decided to examine this kinematic waveform, as this would provide the highest statistical power for the MLE process. Note, linear speed data was used to extract the timing that would separate the start and end time of a forward-deliberate motion (shown in red) and of a backward-spontaneous motion (shown in blue arrow). This was done by finding the timepoint when instantaneous zero linear speed occurs, since this indicates the moment the index finger reaches target. **(B)** Maximum likelihood estimated values for the corresponding histogram on top of each graph. The horizontal axis contains the value of the gradient at the end of the optimization process, and the vertical axis contains the maximum likelihood estimation (MLE) value for the Gamma, normal, exponential and lognormal distributions. Overall, we found that the Gamma and lognormal distributions have a good fit to these kinematics data. However, because Gamma distributions have shown to be a better fit to the kinematics data from individuals with neurological disorders than lognormal distributions, for consistency, we chose to use the Gamma probability distribution for fitting purpose.

**Figure A3.**
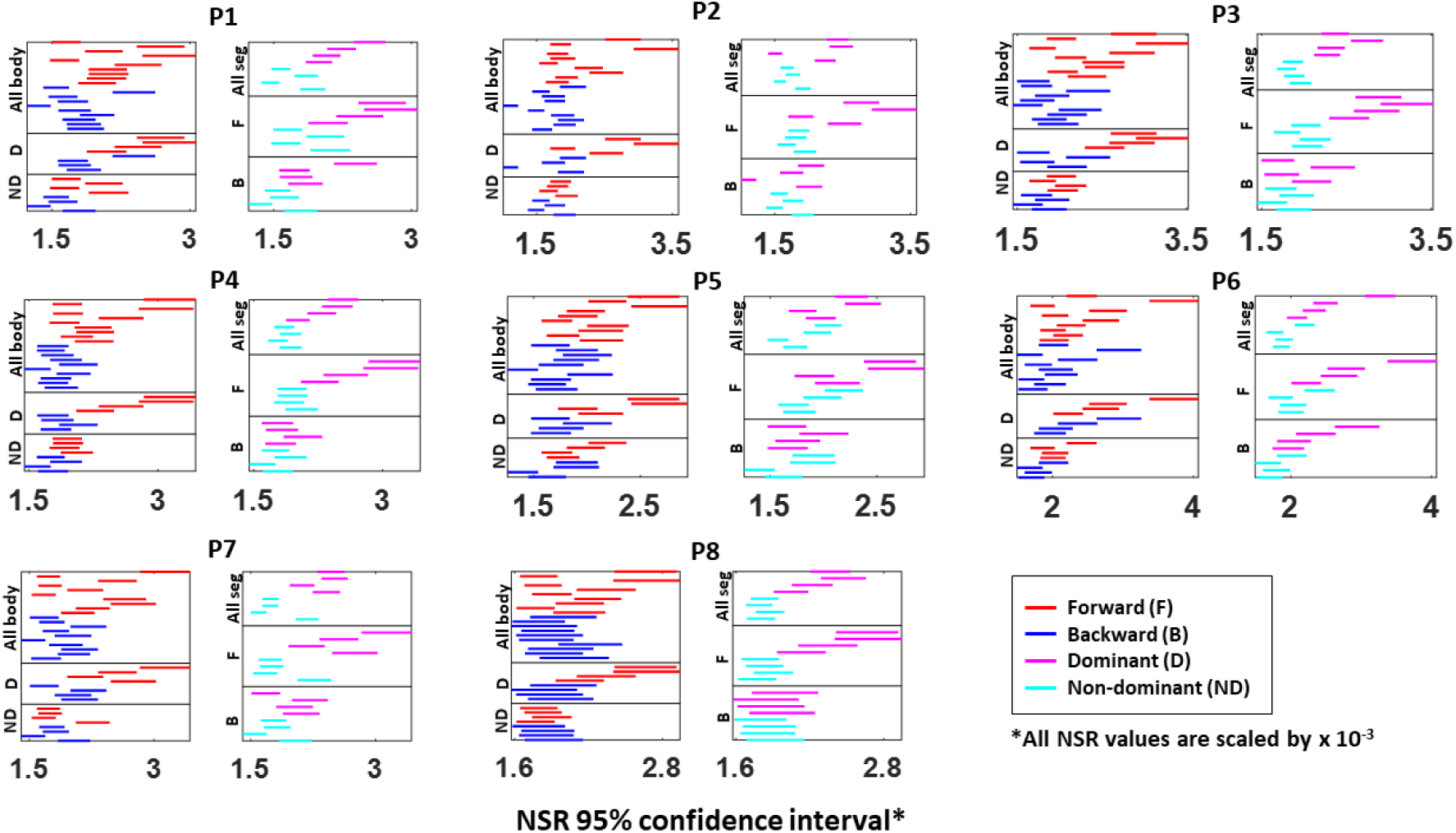
Fitted Gamma scale parameter (i.e., NSR) 95% confidence interval for a single body part’s kinematics data. The 95% confidence interval is plotted for all eight participants (P1 to P8). Each row represents a single body part: under the “All body” category shows all 10 body parts during forward (red) and backward (blue) motions; under the “D (dominant)” category shows the 4 body parts from the dominant side of the arm; under the “ND (non-dominant)” category shows the 4 body parts from the non-dominant side of the arm; under the “All seg (all segment)” category shows the 4 body parts on the dominant (pink) and non-dominant (cyan) side during the entire pointing motion; under the “F (forward)” category shows the 4 body parts on both D and ND side during forward motion; and under the “B (backward)” category shows the 4 body parts on both D and ND side during backward motion.

**Figure A4.**
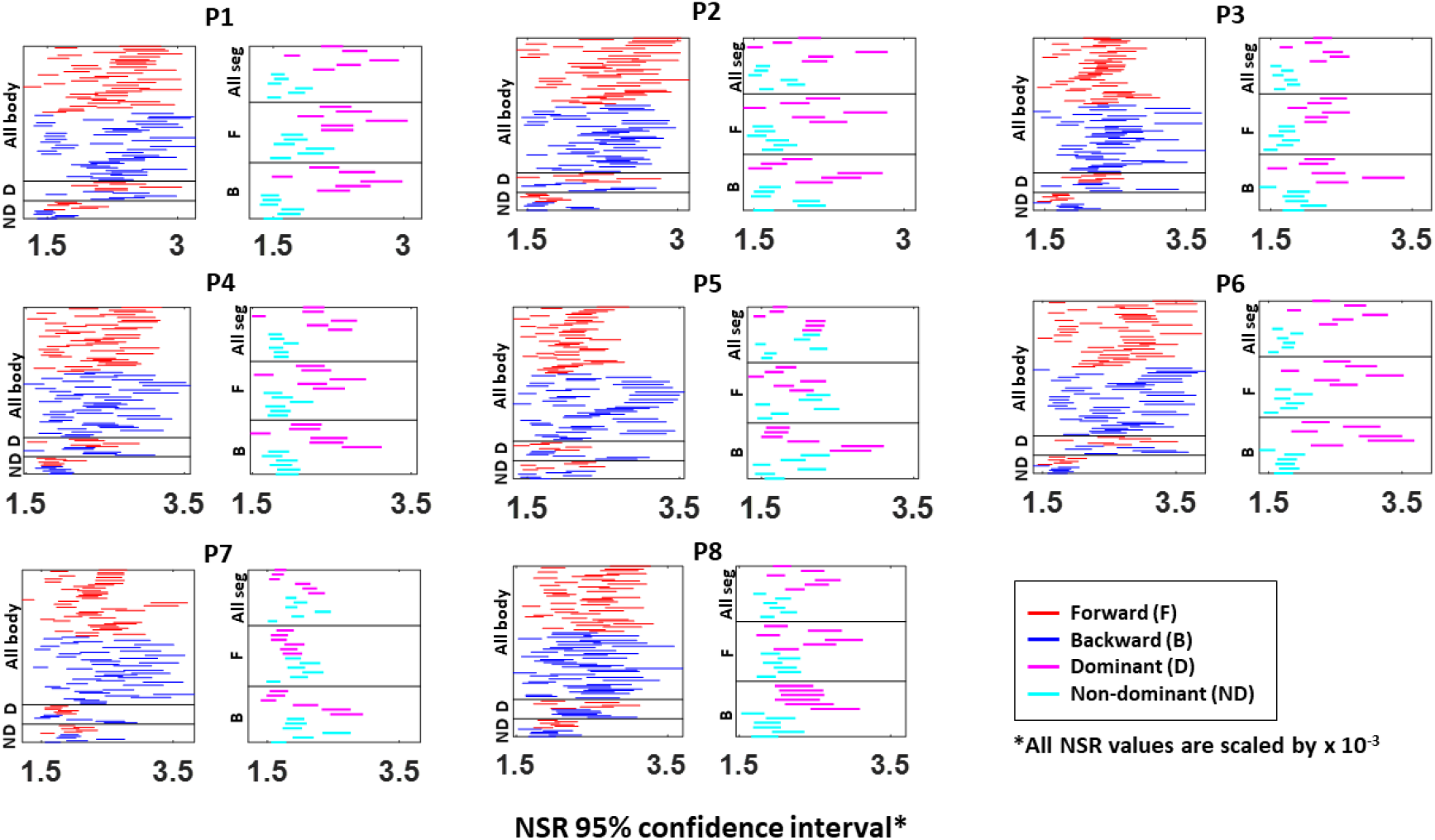
Fitted Gamma scale parameter (i.e., NSR) 95% confidence interval from the absolute difference in kinematics between pairs of body parts. The 95% confidence interval is plotted for all eight participants (P1 to P8). Each row represents a pair of body part: under the “All body” category shows all 45 body part (_10_C_2_) pairs during forward (red) and backward (blue) motions; under the “D (dominant)” category shows the 6 body part pairs (_4_C_2_) from the dominant side of the arm; under the “ND (non-dominant)” category shows the 6 body parts pairs (_4_C_2_) from the non-dominant side of the arm; under the “All seg (all segment)” category shows the 6 body parts pairs (_4_C_2_) on the dominant (pink) and non-dominant (cyan) side during the entire pointing motion; under the “F (forward)” category shows the 6 body parts pairs (_4_C_2_) on both D and ND side during forward motion; and under the “B (backward)” category shows the 4 body parts on both D and ND side during backward motion.

**Figure A5.**
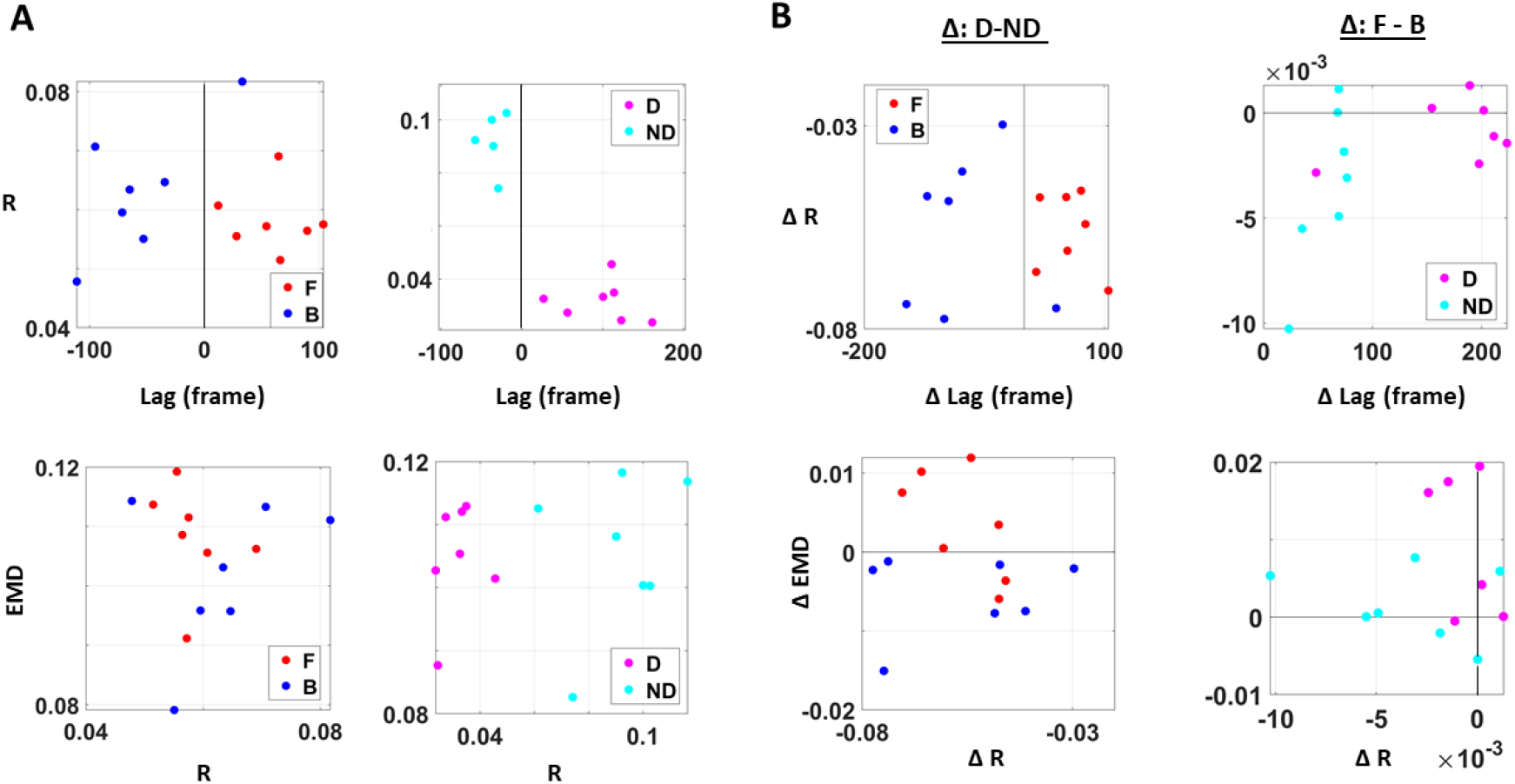
Different viewpoints of the 3D graphs in Figure 8. **(A)** Different viewpoint of graphs in Figure 8B. **(B)** Different viewpoints of graphs in Figure 8C (left) and Figure 8D (right).

**Table A1.** Kolmogorov-Smirnov test statistics (KS-stat) and their p-values (p) on cluster coefficients comparison (left) and cross-correlation (right) between different movement segments (forward (F) vs. backward (B)) and dominance side (dominant (D) vs. non-dominant (ND))1

| Cluster Coefficient |  |  |  |  | Cross-Correlation |  |  |  |  |
| --- | --- | --- | --- | --- | --- | --- | --- | --- | --- |
| Subject | F vs. B |  | D vs. ND |  | Subject | F vs. B |  | D vs. ND |  |
| ID | KS-stat | p | KS-stat | p | ID | KS-stat | p | KS-stat | p |
| P01 | 0.29 | <0.01** | 0.16 | <0.01** | P01 | 0.14 | <0.01** | 0.40 | <0.01** |
| P02 | 0.55 | <0.01** | 0.57 | <0.01** | P02 | 0.55 | <0.01** | 0.57 | <0.01** |
| P03 | 0.38 | <0.01** | 0.16 | <0.01** | P03 | 0.38 | <0.01** | 0.16 | <0.01** |
| P04 | 0.09 | <0.01** | 0.09 | <0.01** | P04 | 0.09 | <0.01** | 0.09 | <0.01** |
| P05 | 0.14 | <0.01** | 0.26 | <0.01** | P05 | 0.14 | <0.01** | 0.26 | <0.01** |
| P06 | 0.17 | <0.01** | 0.28 | <0.01** | P06 | 0.17 | <0.01** | 0.28 | <0.01** |
| P07 | 0.35 | <0.01** | 0.41 | <0.01** | P07 | 0.35 | <0.01** | 0.41 | <0.01** |
| P08 | 0.13 | <0.01** | 0.35 | <0.01** | P08 | 0.13 | <0.01** | 0.35 | <0.01** |

<sup>1</sup>Note, the Kolmogorov-Smirnov test was used, as this test is appropriate for data that do not follow a Gaussian distribution and has a large sample size (n>1000) that may yield low statistical power.

## References

1. Maturana, H.R. and F.J. Varela, Autopoiesis and cognition: The realization of the living. Vol. 42. 1991: Springer Science & Business Media.

2. Abrahamson, D. and R. Sanchez-Garcia, Learning Is Moving in New Ways: The Ecological Dynamics of Mathematics Education. Journal of the Learning Sciences, 2016. 25(2): p. 203–239.

3. Newen, A., The Embodied Self, the Pattern Theory of Self, and the Predictive Mind. Front Psychol, 2018. 9: p. 2270.

4. David, N., A. Newen, and K. Vogeley, The “sense of agency” and its underlying cognitive and neural mechanisms. Conscious Cogn, 2008. 17(2): p. 523–34.

5. Synofzik, M., G. Vosgerau, and A. Newen, I move, therefore I am: a new theoretical framework to investigate agency and ownership. Conscious Cogn, 2008. 17(2): p. 411–24.

6. Frith, C.D., Action, agency and responsibility. Neuropsychologia, 2014. 55: p. 137–42.

7. Tsakiris, M., G. Prabhu, and P. Haggard, Having a body versus moving your body: How agency structures body-ownership. Conscious Cogn, 2006. 15(2): p. 423–32.

8. Tsakiris, M., et al., Neural signatures of body ownership: a sensory network for bodily self-consciousness. Cereb Cortex, 2007. 17(10): p. 2235–44.

9. Toni, I., D. Thoenissen, and K. Zilles, Movement preparation and motor intention. Neuroimage, 2001. 14(1 Pt 2): p. S110–7.

10. Haggard, P. and A. Wing, On the hand transport component of prehensile movements. J Mot Behav, 1997. 29(3): p. 282–7.

11. Torres, E.B., Two classes of movements in motor control. Exp Brain Res, 2011. 215(3-4): p. 269–83.

12. Torres, E.B., Signatures of movement variability anticipate hand speed according to levels of intent. Behav Brain Funct, 2013. 9: p. 10.

13. Ryu, J. and E.B. Torres, Characterization of Sensory-Motor Behavior Under Cognitive Load Using a New Statistical Platform for Studies of Embodied Cognition. Front Hum Neurosci, 2018. 12: p. 116.

14. Torres, E.B., et al., Autism: the micro-movement perspective. Front Integr Neurosci, 2013. 7: p. 2.

15. Bernstein, N.A., The Coordination and Regulation of Movements. 1967, London: Pergamon Press.

16. Torres, E.B., Zipser, D., Reaching to Grasp with a Multi-jointed Arm (I): A Computational Model. Journal of Neurophysiology, 2002. 88: p. 1–13.

17. Scholz, J., Schoner, G., The uncontrolled manifold concept: Identifying control variables for a functional task. Experimental Brain Research, 1999. 126: p. 289–306.

18. Latash, M.L., J.P. Scholz, and G. Schoner, Motor control strategies revealed in the structure of motor variability. Exerc Sport Sci Rev, 2002. 30(1): p. 26–31.

19. Von Holst E. and M. H., The principle of reafference: Interactions between the central nervous system and the peripheral organs., in Perceptual Processing: Stimulus equivalence and pattern recognitione. Dodwell PC, Editor. 1950, Appleton-Century-Crofts: New York. p. 41–72.

20. Wolpert, D.M. and M. Kawato, Multiple paired forward and inverse models for motor control. Neural Netw, 1998. 11(7-8): p. 1317–29.

21. Kawato, M. and D. Wolpert, Internal models for motor control. Novartis Found Symp, 1998. 218: p. 291–304; discussion 304-7.

22. Torres, E.B., Theoretical Framework for the Study of Sensori-motor Integration., in Cognitive Science. 2001, University of California, San Diego: La Jolla. p. 115.

23. Nishikawa, K., S. Murray, and M. Flanders, Do arm postures vary with the speed of reaching? Journal of Neurophysiology, 1999. 81(5): p. 2582–6.

24. Atkeson, C.G. and J.M. Hollerbach, Kinematic features of unrestrained vertical arm movements. J Neurosci, 1985. 5(9): p. 2318–30.

25. Torres, E.B., Zipser D., Simultaneous control of hand displacements and rotations in orientation-matching experiments. Journal of Applied Physiology, 2004. 96(Highlighted Topic Neural Control of Movement): p. 1978–1987.

26. Torres, E.B., New symmetry of intended curved reaches. Behav Brain Funct, 2010. 6: p. 21.

27. Torres, E. and R. Andersen, Space-time separation during obstacle-avoidance learning in monkeys. J Neurophysiol, 2006. 96(5): p. 2613–32.

28. Kalampratsidou, V. and E.B. Torres, Peripheral Network Connectivity Analyses for the Real-Time Tracking of Coupled Bodies in Motion. Sensors (Basel), 2018. 18(9).

29. Torres, E.B., K.M. Heilman, and H. Poizner, Impaired endogenously evoked automated reaching in Parkinson’s disease. J Neurosci, 2011. 31(49): p. 17848–63.

30. Yanovich, P., et al., Spatial-orientation priming impedes rather than facilitates the spontaneous control of hand-retraction speeds in patients with Parkinson’s disease. PLoS One, 2013. 8(7): p. e66757.

31. Torres, E.B., P. Yanovich, and D.N. Metaxas, Give spontaneity and self-discovery a chance in ASD: spontaneous peripheral limb variability as a proxy to evoke centrally driven intentional acts. Front Integr Neurosci, 2013. 7: p. 46.

32. Torres, E.B., et al., Toward Precision Psychiatry: Statistical Platform for the Personalized Characterization of Natural Behaviors. Front Neurol, 2016. 7: p. 8.

33. Torres, E.B., Atypical signatures of motor variability found in an individual with ASD. Neurocase, 2011. 19(2): p. 150–65.

34. Mosimann, J.E., Size allometry: size and shape variables with characterizations of the lognormal and generalized gamma distributions.. J. Am. Stat. Assoc., 1970. 65: p. 930–945.

35. Lleonart, J., J. Salat, and G.J. Torres, Removing allometric effects of body size in morphological analysis. J Theor Biol, 2000. 205(1): p. 85–93.

36. Monge, G., Memoire sur la theorie des deblais et des remblais., in Histoire de ? Academie Royale des Science; avec les Memoired de Mathematique et de Physique. 1781, De L’imprimerie Royale: Paris, France.

37. Arjovsky, M., S. Chintala, and L. Bottou. Wasserstein Generative Adversarial Networks. in Proceedings of the 34th International Conference on Machine Learning. 2017. Sydney, Australia.

38. Stolfi, J. and L.J. Guibas, The earth mover’s distance as a metric for image retrieval.. Int. J. Comput. Vis., 2000. 40: p. 99–121.

39. Rubner, Y., L.J. Guibas, and C. Tomasi. The earth mover’s distance, multi-dimensional scaling, and color-based image retrieval. in Proceedings of the ARPA image understanding workshop. 1997.

40. McClelland, J. and D. Koslicki, EMDUniFrac: exact linear time computation of the unifrac metric and identification of differentially abundant organisms. Journal of mathematical biology, 2018. 77(4): p. 935–949.

